# ST8SIA4-mediated polysialylation is critical for CCR2-driven monocyte egress from the bone marrow

**DOI:** 10.64898/2026.01.09.698551

**Authors:** Aurélien Boyancé, Nelly Gilles, Maxime Rotival, Alexandre Stella, Camille Gorlt, Karima Chaoui, Sarah Monard, Nacho Aguiló, Thomas Mangeat, Odile Burlet-Schiltz, Hugues Lortat-Jacob, Anja Münster-Kühnel, Anne Gonzalez de Peredo, Denis Hudrisier, Lluis Quintana-Murci, Rabia Sadir, Herbert Hildebrandt, Olivier Neyrolles, Yoann Rombouts

## Abstract

Polysialylation is a rare post-translational protein modification essential for brain development and synaptic plasticity, where it fine-tunes cell–cell and cell–matrix interactions to direct neuronal migration, neurite outgrowth and synaptogenesis. Although polysialic acid is also expressed on circulating leukocytes, its functions in the immune system remain largely unexplored. Guided by analysis of publicly available human genomic data showing that naturally occurring variants of *ST8SIA4*, which encodes one of the two enzymes that mediate polysialylation, associate with reduced circulating monocyte counts, we investigated the *in vivo* role of this enzyme in monocyte biology. Using *St8sia4*-deficient mice, we show that ST8SIA4-dependent polysialylation is essential for CCR2-mediated egress of inflammatory monocytes from the bone marrow at steady state and during *Mycobacterium tuberculosis* infection. We confirm NCAM1 as the principal polysialylated protein in inflammatory monocytes and demonstrate that *Ncam1*-deficient mice phenocopy the monocyte defects observed in *St8sia4*-deficient animals. Mechanistically, loss of ST8SIA4-dependent polysialylation impairs engagement and internalization of the CCR2 ligands CCL2 and CCL7, accompanied by disrupted CCR2 surface organization and extensive cytoskeletal remodeling. Together, these findings identify polysialylation as a previously unrecognized regulator of CCR2 function and monocyte mobilization, with broad implications for immune surveillance and inflammatory responses.

**Highlights:**

- Human *ST8SIA4* variants are associated with reduced blood monocyte and lymphocyte counts and increased risk of SLE.
- *St8sia4*^-/-^ mice exhibit monocytopenia with impaired CCR2-mediated egress of inflammatory monocytes from the bone marrow both at steady state and during *Mycobacterium tuberculosis* infection.
- ST8SIA4 polysialylates NCAM1 (CD56) in inflammatory monocytes, and *Ncam1*^-/-^
- mice phenocopy the monocytopenia observed in *St8sia4*^-/-^ mice.
- Loss of ST8SIA4 and polysialylation impairs CCR2-mediated binding and endocytosis of CCL2 and CCL7 by monocytes.
- Polysialic acid itself does not function as a co-receptor for these chemokines, revealing a new paradigm for its role in immune cell trafficking.
- Proper CCR2 surface distribution and function, as well as cytoskeletal organization in inflammatory monocytes, depend on ST8SIA4-mediated polysialylation.

## Introduction

Glycans and lectins play key roles in directing leukocyte trafficking between the blood, lymphatics, and peripheral tissues, a process fundamental to the initiation, maintenance, and regulation of immune responses. For example, leukocyte adhesion to the vascular endothelium relies on the binding of selectins to specific glycan epitopes, such as sialyl Lewis X and its variants, that decorate the extracellular domains of membrane glycoproteins including CD44 and PSGL-1.^1^ Likewise, the migration of immune cells into inflamed tissues along chemokine gradients is shaped by glycosaminoglycans (GAGs)—negatively charged sulfated and carboxyl group-containing carbohydrate polymers that bind chemokines with high affinity.^2^ GAG-chemokine interactions immobilize chemokines on endothelial proteoglycans and protect them from proteolysis, thereby ensuring that leukocyte recruitment occurs at spatially restricted sites. The extensive structural diversity of GAGs, together with the differential affinities of individual chemokines for these carbohydrate polymers contributes to the remarkable specificity of chemokine-guided leukocyte trafficking.

Polysialic acid (polySia) is another negatively charged polysaccharide, composed of a linear homopolymer of a2,8-linked sialic acid residues. In mammals, polySia is synthesized in the Golgi apparatus by the polysialyltransferases ST8SIA2 and ST8SIA4, which add this modification to *N*- and *O*-linked glycans on select glycoproteins. To date, roughly a dozen polysialylated proteins have been identified, including the neural cell adhesion molecule 1 (NCAM1/CD56), neuropilin-2 (NRP2), C−C chemokine receptor-7 (CCR7), and ST8SIA2 and ST8SIA4 themselves.^3^ PolySia is highly enriched in the central nervous system (CNS), where it regulates tissue development, cell migration, axon guidance, and synaptic plasticity.^3,4^ Outside the CNS, polySia is expressed by several immune cell types in both humans and mice, including monocytes, macrophages, and dendritic cells (DCs),^5–10^ as well as neutrophils (in mice) and natural killer cells and activated T cells (in humans).^11,12^ In immune cells, polySia is produced predominantly by ST8SIA4.

Most studies investigating the role of polysialylation in immune regulation have relied on exogenous polySia to stimulate immune cells. Across both *in vitro* and *in vivo* settings, these studies consistently demonstrate that this carbohydrate homopolymer exerts potent anti-inflammatory effects.^13–18^ In particular, exogenous polySia engages inhibitory sialic-acid–binding immunoglobulin-like lectin (Siglec) receptors, dampening cellular maturation and suppressing the production of pro-inflammatory cytokines.

A second line of studies has focused on cells or mice lacking the polysialyltransferase ST8SIA4 (*St8sia4^-/-^*) to investigate how endogenous polysialylation regulates immune cell function. Drawing on parallels from the CNS, polySia is often considered an anti-adhesive molecule that disrupts cell–cell interactions, thereby impairing macrophage phagocytosis and reducing neutrophil adhesion to the vascular endothelium during extravasation.^6,19^ Beyond modulating direct cell–cell contacts, cell-surface polySia also influences interactions between immune cells and soluble proteins. In human and mouse DCs, polySia carried by NRP2 and/or CCR7 functions as a co-receptor for the chemokine CCL21, facilitating CCR7-dependent migration of DCs from tissues into lymphatic vessels.^8,10,20,21^ In neutrophils, by contrast, loss of polySia enhances binding of chemokines, such as CXCL1 and N-formylmethionyl-leucyl-phenylalanine, yet paradoxically diminishes chemotaxis, likely through effects on chemokine-receptor signaling as well as migratory velocity and persistence.^19^ Polysialylation has also been implicated in regulating bone-marrow egress of hematopoietic progenitors and their entry into the thymus,^22^ although whether these effects are mediated primarily through altered cell–cell interactions, chemokine engagement, or other mechanisms remains unresolved.

Taken together, these findings indicate that polySia modulates immune function in a highly cell- and protein-specific manner. Elucidating the underlying mechanisms promises to reveal fundamental principles immune-cell regulation in both homeostatic and pathological contexts.

In this study, we investigated the role of polysialylation in monocytes. Analysis of publicly available human genomic data revealed that variants in *ST8SIA4* are associated with reduced blood monocyte and lymphocyte counts. Using *St8sia4^-/-^*mice, we show that ST8SIA4 is critical for monocyte homeostasis, with inflammatory monocyte numbers diminished both at steady-state and during *Mycobacterium tuberculosis* infection. Through mixed bone-marrow chimera experiments and *in vitro* and *in vivo* chemotaxis assays, we demonstrate that this phenotype reflects cell-intrinsic defects in polySia-deficient inflammatory monocytes, which fail to efficiently bind and internalize the chemokines CCL2 and CCL7, leading to impaired bone marrow egress. We confirm that NCAM1 (CD56) is the principal polySia carrier in inflammatory monocytes, and show that *Ncam1^-/-^* mice display a similar reduction in circulating inflammatory monocytes. Importantly, polySia does not function as a co-receptor for CCL2 or CCL7; rather its absence disrupts cytoskeletal architecture, altering CCR2 localization and function. Together, our findings reveal a previously unrecognized role for polysialylation in sustaining CCR2 activity and promoting inflammatory monocyte mobilization during homeostasis and infection.

## Results

### *ST8SIA4* variants associate with decreased monocyte and lymphocyte counts and systemic lupus erythematosus in humans

To investigate the function of *ST8SIA4* in humans, we searched the Open Targets Platform^23,24^ for phenotypes associated with the *ST8SIA4* locus, and extracted 95% credible sets of fine-mapped variants associated with each phenotype (such that each set has >95% probability to contain the causal variant for the association). We found that *ST8SIA4* variants are associated with both lymphocyte and monocyte counts as well as systemic lupus erythematosus (SLE) (Figure 1A, index single-nucleotide polymorphisms (SNPs) rs2725117-A>C & rs2548257-C>A, and rs12153670-A>G, respectively, Figure S1A). Index SNPs for leukocyte count phenotypes were in strong linkage disequilibrium (LD) with each other (r^2^=0.86), with the same allele decreasing counts for both lymphocytes and monocytes; however, both phenotypes had fully disjoint credible sets, suggesting distinct causal variants (Figure S1B). The lead SLE variant, rs12153670-A>G, displayed only moderate LD with index SNPs for leukocyte count phenotypes (r^2^=0.43-0.5), with risk alleles for SLE being associated with lower monocyte/lymphocyte counts. While there was a significant overlap between the 95% credible set of SLE and monocyte counts, we found limited evidence for shared causal variants (colocalization probability P_H4_=0.6). Altogether, this suggests that distinct genetic variants at the *ST8SIA4* locus separately influence leukocyte cell counts and SLE risk in humans.

**Figure 1:**
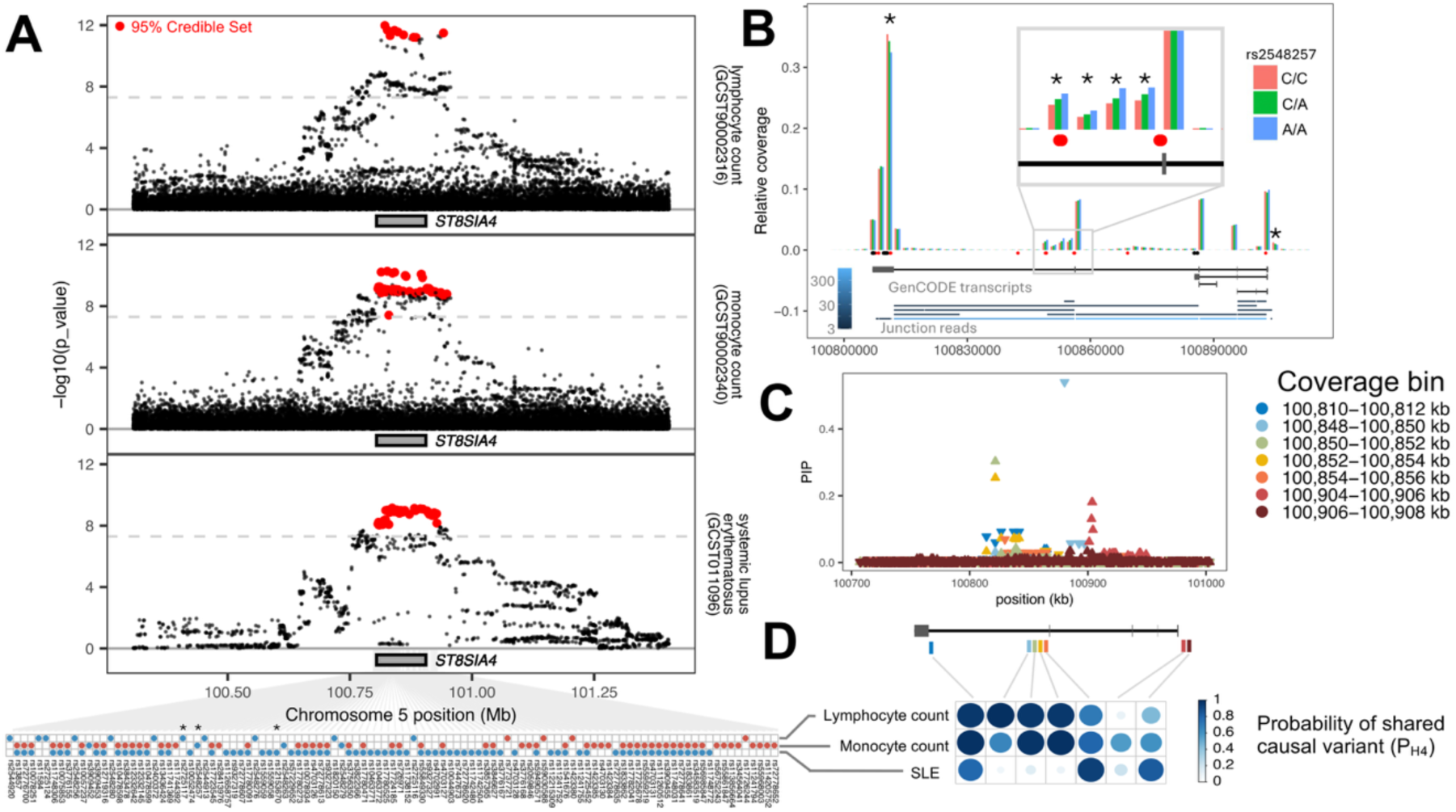
*ST8SIA4* variants and isoform usage associate with decreased monocyte and lymphocyte counts and systemic lupus erythematosus. (A) Significant GWAS hits at the *ST8SIA4* locus. -log_10_(association *p*-value) are shown for all tested variants within 500 kb of the ST8SIA4 gene (grey rectangle). For each phenotype, we highlight in red the set of variants with >95% probability to contain the causal variants (95% credible set). Bottom panel shows the overlap of 95% credible sets for the 3 phenotypes, with red and blue respectively indicating positive/negative effect of the alternative allele on the phenotype. Stars indicate the index SNPs associated with each phenotype. (B) Normalized coverage of *ST8SIA4* in resting CD14^+^ monocytes, according to the genotype of the variant most strongly associated with monocytopenia, rs2548257-C>A). Coverage is computed in bins of 2 kb for visualization purposes. Structure of GenCODE *ST8SIA4* transcripts is shown in black, and observed junctions are shown in shades of blue (lighter for higher coverage). Black and red dots annotate polyA sites inferred from either micro-well seq experiments (scUTRquant UTRome) and or 3’ Region Extraction and Deep Sequencing (3’READS, polyA DB v3.2). (*) significant QTL at FDR<1%. (C) Fine mapping of *ST8SIA4* coverage QTLs. For each genomic bin with a significant coverage QTL, we display the probability of each variant to causally impact read coverage (posterior inclusion probabilities, or PIP) as a function of variant location. (D) Colocalization results showing the posterior probability of a shared causal variant to impact both coverage in a specific genomic bin and phenotypic traits (P_H4_).

### Altered isoform usage drives effects of *ST8SIA4* variants on leukocytopenia and systemic lupus erythematosus

To investigate the molecular basis of these associations, we first assessed the colocalization of GWAS hits with molecular quantitative trait loci (QTLs) from the eQTL catalog,^25^ which revealed that variants associated with both SLE and monocyte counts, but not lymphocyte counts, colocalize with variants altering isoform usage of *ST8SIA4* across multiple immune cell types, including both resting and stimulated monocytes (Student’s P<2.2x 10^-4^, P_H4_ >0.8) (Table S1A). Because colocalizing QTLs were associated with a diversity of alternative splicing events, we analyzed *ST8SIA4* coverage in bulk RNAseq of resting CD14^+^ monocytes to resolve the impact of monocytopenia-associated variants on *ST8SIA4.*^26^ Our analysis revealed that the monocytopenia-associated allele rs2548257-A correlates with a decrease in coverage at the major polyadenylation site on exon 5 (R^2^=0.24, student’s P< 1.4 x 10^-13^, P_H4_ = 0.98) together with a strong increase in coverage of intron 5 (max R^2^ across tested bins = 35%, student’s P< 3.8 x 10^-9^, max P_H4_ = 0.98), due to the presence of several unannotated polyadenylation sites within 8 kb from exon 4 (Figure 1B and tables S1B-C).^27^ Interestingly, we found that coverage of intron 4 was under the control of several distinct genetic variants, as we moved further away from exon 4 (Figure 1C). In particular, SLE risk variants specifically colocalized with variants that control the 2 kb located immediately downstream of exon 4, (P_H4_ > 0.93), whereas variants associated with decreased blood monocyte and lymphocyte counts preferentially colocalized with regions located further downstream (max P_H4_ > 0.99 in both cases, Figure 1D). In addition, SLE variants further colocalized with promoter variants associated with upstream TSS usage (P_H4_ > 0.83, Figure 1D). Altogether, these data indicate that variants linked to leukocyte count and SLE promote increased usage of alternative polyadenylation sites, likely resulting in premature termination of *ST8SIA4* transcription and reduced expression of the full-length, canonical isoform of the enzyme.

### ST8SIA4 deficiency causes monocytopenia in mice

In order to assess the role of ST8SIA4 and polySia in the regulation of leukocyte homeostasis *in vivo*, we analyzed leukocyte populations in wild-type (WT) and *St8sia4^- /-^* mice.^28^ We detected strong polySia expression on the surface of blood-derived myeloid cells (CD45^+^ CD11b^+^), but not on lymphoid cells (CD45^+^ CD11b^-^), by flow cytometry (Figures 2A and S2A). Among myeloid cells, polySia was highly expressed by inflammatory monocytes (CD45^+^, CD11b^+^, Ly6C^+^, CD115^+^), eosinophils (CD45^+^, CD11b^+^, Siglec-F^+^) and neutrophils (CD45^+^, CD11b^+^, Ly6G^+^) but not by patrolling monocytes (CD45^+^, CD11b^+^, Ly6C^low^, CD115^+^) (Figure 2B, S2A-B). As expected, polySia expression was absent in ST8SIA4-deficient myeloid cells, confirming that ST8SIA4 is the main enzyme responsible for polysialylation in these cells (Figures 2A-B).

**Figure 2:**
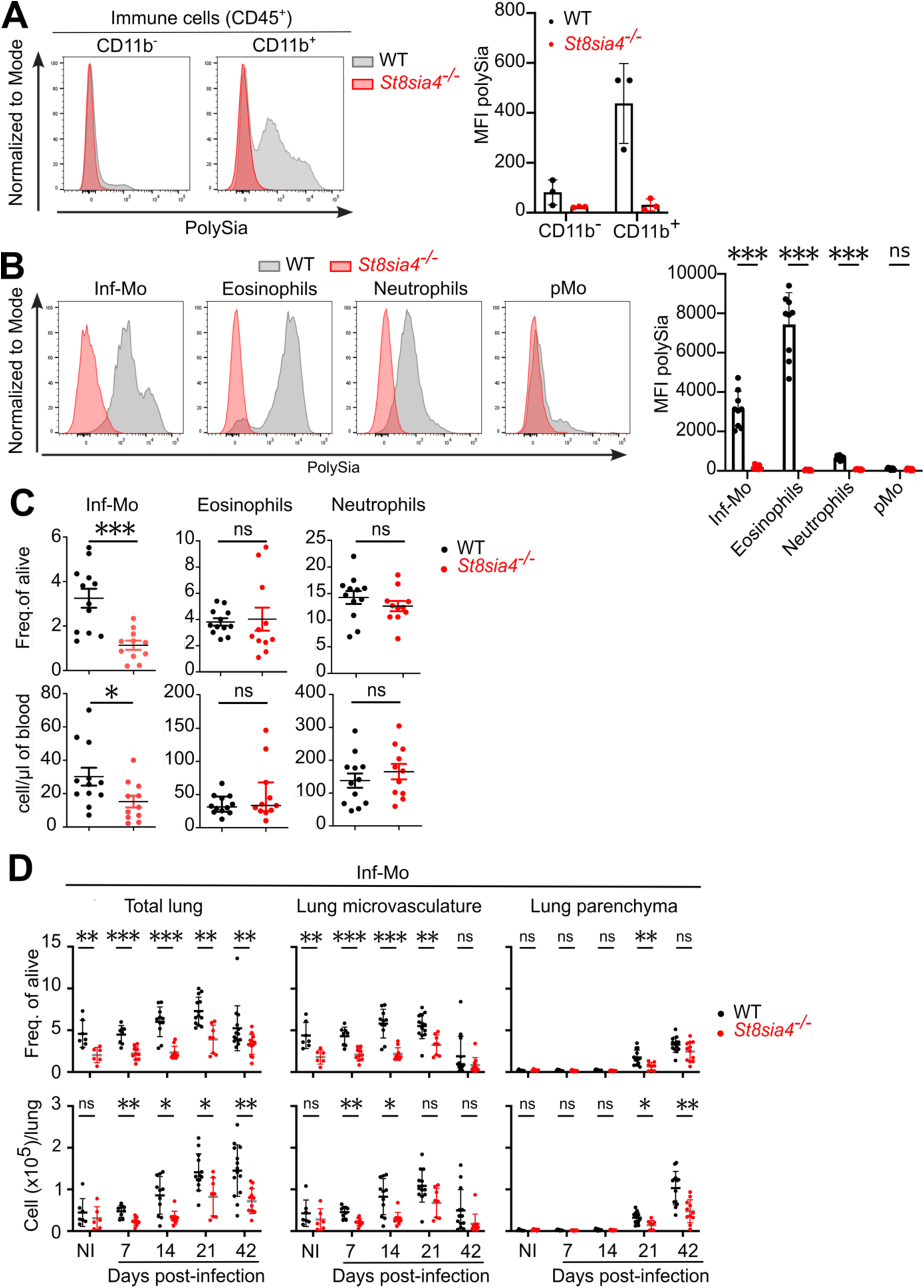
*St8sia4* deficiency in mouse causes monocytopenia. (A) Representative flow cytometry histograms (left panel) showing polySia surface expression on blood lymphoid (CD45⁺ CD11b⁻) and myeloid (CD45⁺ CD11b⁺) cells from WT and *St8sia4^-/-^*mice, with corresponding geometric mean fluorescence intensity (MFI) values summarized in the bar plot (right panel). Each MFI data point corresponds to an individual mouse (n=3 mice per group). Bars and error indicate the mean ± standard deviation. (B) Similar to A for circulating myeloid cell subpopulations including inflammatory monocytes (Inf-Mo; CD45^+^, CD11b^+^, Ly6C^+^, CD115^+^), eosinophils (CD45^+^, CD11b^+^, siglec-F^+^), neutrophils (CD45^+^, CD11b^+^, Ly6G^+^) and patrolling monocytes (pMo; CD45^+^, CD11b^+^, Ly6C^low^, CD115+). Each MFI data point corresponds to an individual mouse. Bars and error indicate the mean ± standard deviation (n=7-9 mice per group, pooled from 3 independent experiments). Statistical analysis was performed using Mann-Whitney *t*-tests corrected for multiple comparison using Holm-Šídák method. *** indicates an adjusted *p*-value ≤0.001. The abbreviation ’ns’ stands for non-significant. (C) Flow cytometry measurement of the frequency (top) and number (bottom) of circulating inflammatory monocytes (Inf-Mo), eosinophils and neutrophils in WT and *St8sia4^-/-^* mice (n=11-12 mice per group, pooled from 4 independent experiments). Each data point corresponds to an individual mouse. Bars and error indicate the mean ± standard error of the mean. Statistical analysis was performed using Wech’s *t*-tests. * and *** indicate *p*-values ≤0.05 and ≤0.001 respectively. The abbreviation ’ns’ stands for non-significant. (D) Flow cytometry measurement of the frequency (top) and number (bottom) of inflammatory monocytes (Inf-Mo) in total lung, microvasculature and parenchyma in non-infected (NI) and during infection of WT and *St8sia4^-/-^* mice with *Mycobacterium tuberculosis*. Each data point corresponds to an individual mouse (n=6-14 mice per group per time point, pooled from at least 2 independent experiments per time point). Bars and error indicate the mean ± standard deviation. Statistical analysis was performed using Mann-Whitney *t*-tests corrected for multiple comparison using Holm-Šídák method. *, **, *** indicate adjusted *p*-values ≤0.05, ≤0.01 and ≤0.001 respectively. The abbreviation ’ns’ stands for non-significant.

We next asked whether the absence of polySia expression affects the abundance of these cells in the blood. Whereas WT and *St8sia4^-/-^* mice exhibited similar percentages and absolute numbers of circulating neutrophils and eosinophils, there was a marked reduction in inflammatory monocytes in *St8sia4^-/-^* mice (Figure 2C).

To assess whether this monocytopenia impairs the recruitment of inflammatory monocytes to peripheral tissues, we intranasally infected mice with *Mycobacterium tuberculosis* (*Mtb*) H37Rv, a model that specifically recruits monocytes to the lungs without triggering emergency myelopoiesis.^29^ We observed a significant decrease in the percentage and absolute number of inflammatory monocytes in the lung of *St8sia4^- /-^* mice as early as 7 days post-infection (Figures 2D and S2B). To determine whether these inflammatory monocytes were located in the pulmonary vasculature or the parenchyma, we labeled circulating immune cells by intravenously injecting an anti-CD45.2 antibody shortly before sacrificing the mice.^30^ *St8sia4^-/-^* mice exhibit reduced percentages and numbers of vascular inflammatory monocytes at 7 and 14 days post-infection (Figure 2D). Consistently, infiltration of inflammatory monocytes into the lung parenchyma was decreased at 21 days post-infection (Figure 2D). No notable differences were detected in other innate immune cell populations. However, *St8sia4*^- /-^ mice exhibited reduced percentages and absolute numbers of Th1 (T-bet⁺) and Th17 (RORγt⁺) CD4⁺ T cells at 21 days, but not at 41 days, post-infection (Figures S2B–C and S3A–B). Pulmonary bacterial burden was comparable between WT and *St8sia4*^-/-^mice at days 14 and 21 post-infection, but was modestly reduced in *St8sia4*^-/-^ mice by day 42 (Figure S3C). This phenotype mirrors that of *Ccr2*^-/-^ mice infected with the same *Mtb* strain, which also show reduced recruitment of inflammatory monocytes to the lungs and delayed T-cell responses, yet do not display increased susceptibility to tuberculosis.^31,32^

Taken together, these results demonstrate that *St8sia4^-/-^*mice have reduced numbers of circulating inflammatory monocytes, both at steady state and during *Mtb* infection.

### ST8SIA4 is required for the bone marrow egress of inflammatory monocytes in response to CCL2 and CCL7

We next asked whether the reduced numbers of inflammatory monocytes in the blood and lungs of *St8sia4*^-/-^ mice resulted from defects in their production, survival, mobilization from the BM, or conversion to patrolling monocytes. We found comparable numbers of inflammatory monocytes and their progenitors in the BM of WT and *St8sia4*^-/-^ animals (Figures S4A-B), and concentration of macrophage colony-stimulating factor (M-CSF), a master regulator of myelopoiesis, was also similar in both the serum and BM extracellular fluid of WT and *St8sia4*^-/-^ mice (Figure S4C). Moreover, loss of ST8SIA4 had no detectable impact on the *in vitro* differentiation of granulocyte-monocyte progenitors (GMP) into macrophages, granulocytes, or mixed granulocyte-macrophage colonies (Figure S4D), indicating that *St8sia4*^-/-^ deficiency did not lead to defective monocyte development.

Inflammatory monocytes constitute obligatory precursors of patrolling monocytes at steady-state.^33^ We therefore asked whether exacerbated conversion of inflammatory monocytes into patrolling monocytes might explain the lower numbers of circulating inflammatory monocytes in *St8sia4*^-/-^ mice. However, we found similar numbers of patrolling monocytes in the blood and BM of WT and *St8sia4*^-/-^ mice, ruling out this possibility (Figure S5A).

Given that the spleen serves as a reservoir for inflammatory monocytes and contributes to their rapid deployment into the bloodstream,^34^ we next quantified splenic inflammatory monocytes in WT and *St8sia4*^-/-^ mice. As in blood, we detected about two times fewer inflammatory monocytes in the spleen of *St8sia4*^-/-^ mice than in WT animals (Figure S5B), demonstrating that retention of inflammatory monocytes in the spleen does not explain the reduction in circulating inflammatory monocytes in the absence of *St8sia4*. We also ruled out the possibility of excess cell death of ST8SIA4-deficient inflammatory monocytes, with similar proportions of caspase-activated inflammatory monocytes detected in the blood or BM of WT and *St8sia4*^-/-^ mice (Figure S5C).

Lastly, we asked whether mobilization of inflammatory monocytes from the BM to the peripheral circulation was deficient in *St8sia4*^-/-^ mice. In view of the role of CXCL12-CXCR4 interaction in promoting retention of inflammatory monocytes in the BM and of CCL2 and CCL7 interactions with CCR2 in stimulating egress from the BM,^35–38^ we measured expression of these molecules in WT and *St8sia4*^-/-^ mice. We found no differences in CXCR4 expression on BM monocytes or CXCL12 concentrations in serum and BM extracellular fluid between WT and *St8sia4^-/-^*mice (Figure S5D-E). In contrast, we detected a higher percentage of CCR2^+^ inflammatory monocytes in the BM of *St8sia4*^-/-^ mice compared to WT mice (Figure 3A). We also detected a slight elevation in serum CCL2 levels and a substantial increase CCL7 concentrations in the serum of *St8sia4*^-/-^ mice (Figure 3B).

**Figure 3:**
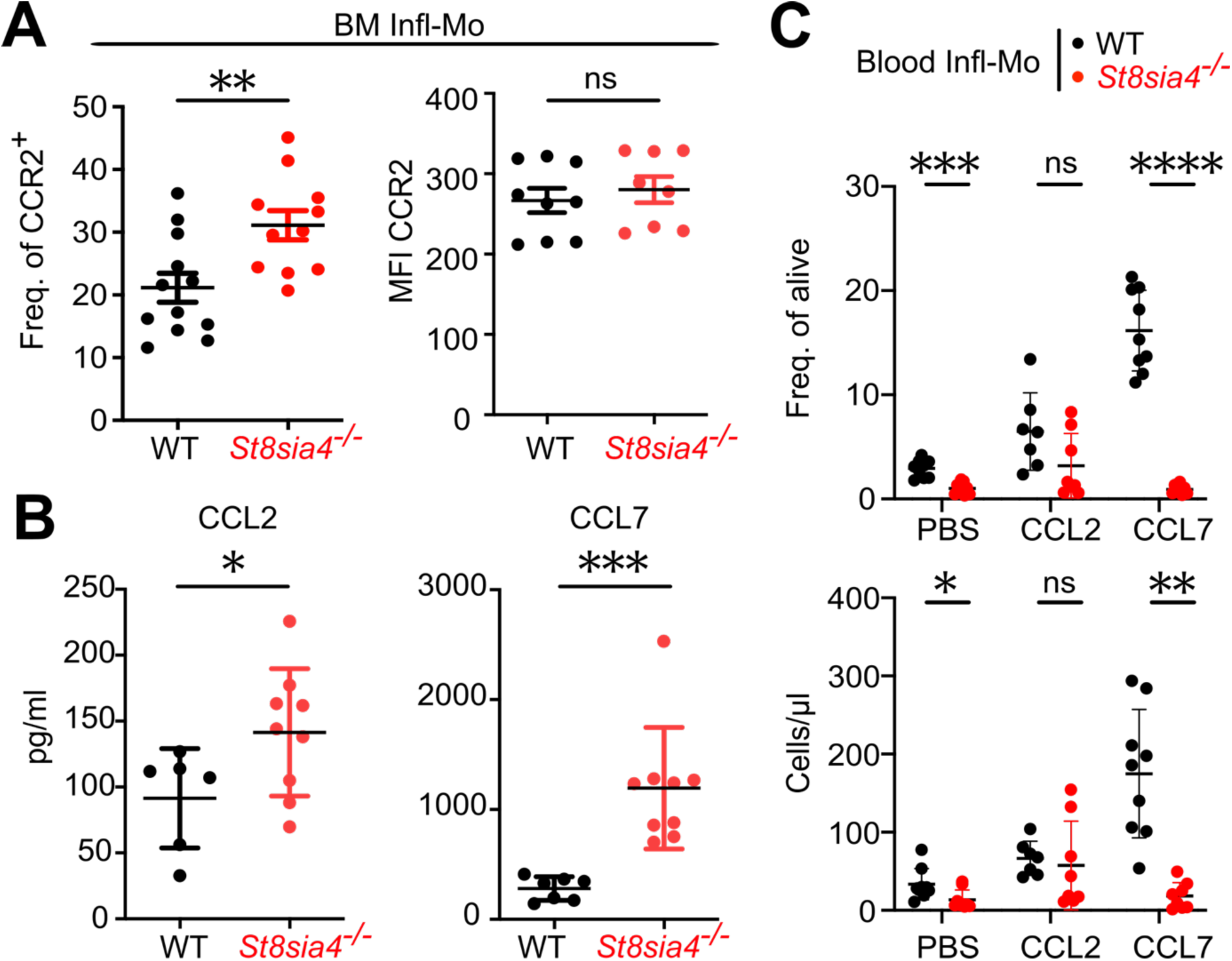
ST8SIA4 is required for bone marrow exit of inflammatory monocytes in response to CCL2 and CCL7. (A) Frequency (left) and representative median fluorescence intensity (MFI; right panel) of CCR2 surface expression on bone marrow (BM) inflammatory monocytes (Inf-Mo) from WT and *St8sia4^-/-^* mice, as measured by flow cytometry. Each data point corresponds to an individual mouse (n=11-12 and 8-9 mice per group for frequency and MFI respectively, pooled from 3-4 independent experiments). Bars and error indicate the mean ± standard error of the mean. Statistical analysis was performed using Wech’s *t*-tests. ** indicates a *p*-value ≤0.01. (B) Concentration of CCL2 and CCL7 in the serum of WT and *St8sia4^-/-^*mice, as measured by ELISA. Each data point corresponds to an individual mouse (n=6-9 and 7-9 mice per group for CCL2 and CCL7 respectively, pooled from 2 independent experiments). Bars and error indicate the mean ± standard deviation. Statistical analysis was performed using Welch’s *t*-tests. * and *** indicate *p*-values ≤0.05 and ≤0.001 respectively. (C) Flow cytometry analysis of the frequency (top) and number (bottom) of circulating inflammatory monocytes in WT and *St8sia4^-/-^* mice at 30 minutes following intravenous injection of vehicle (DPBS), CCL2 or CCL7. Each data point corresponds to an individual mouse (n=7-9 mice per group per condition, pooled from 2 independent experiments). Bars and error indicate the mean ± standard deviation. Statistical analysis was performed using Welch’s *t*-tests corrected for multiple comparison using Holm-Šídák method. *, **, *** and **** indicate adjusted *p*-values ≤0.05, ≤0.01, ≤0.001 and ≤0.0001 respectively.

A previous study showed that mice deficient in CCR1, CCR2, CCR3, and CCR5 (iCCR mice) have higher plasma concentrations of their cognate chemokines (CCL5, CCL7 and CCL11), suggesting that these receptors actively internalize their respective chemotactic ligands under homeostatic conditions.^38^ iCCR-deficient mice also display monocytopenia,^38^ raising the possibility that ST8SIA4-deficient monocytes may similarly have impaired CCR2-mediated chemokine endocytosis.

To test this hypothesis, we injected WT and *St8sia4^-/-^*mice intravenously with CCL2, CCL7, or vehicle, and quantified inflammatory monocyte recruitment into the blood by flow cytometry. As expected, both chemokines triggered the recruitment of inflammatory monocytes in WT mice, with CCL7 having a more potent effect than CCL2 (Figure 3C). *St8sia4^-/-^* mice showed a modest increase in circulating inflammatory monocytes in response to CCL2, although below the level observed in WT mice. In contrast, inflammatory monocyte mobilization in response to CCL7 was nearly completely absent in *St8sia4^-/-^* mice (Figure 3C).

Altogether, these findings indicate that *St8sia4^-/-^*mice exhibit defective egress of inflammatory monocytes from the BM in response to CCL7 and, to a lesser extent, CCL2.

### Defective chemokine binding and endocytosis by monocytes drive monocytopenia in *St8sia4^-/-^* mice

We next asked whether the monocytopenia observed in *St8sia4^-/-^* mice reflects a cell-intrinsic defect in inflammatory monocyte migration. As no model currently allows monocyte-specific deletion of ST8SIA4, we first used mixed bone marrow chimeras to assess whether this phenotype originates within the hematopoietic compartment. Mixed chimeras were generated by reconstituting lethally irradiated WT or *St8sia4^-/-^*mice with a 50:50 mix of BM cells from CD45.1^+^ WT and CD45.2^+^ *St8sia4^-/-^* donor animals (Figures 4A and S5F). In both recipient WT and *St8sia4^-/-^* animals, circulating CD45.1^+^ WT inflammatory monocytes significantly outnumbered CD45.2^+^ ST8SIA4-deficient inflammatory monocytes (Figure 4B). Conversely, we found a slightly higher proportion of inflammatory CD45.2^+^ ST8SIA4-deficient inflammatory monocytes in the BM of reconstituted mice, suggesting a defect in egress of these cells from the BM. As anticipated, the proportions of CD45.1^+^ WT and CD45.2^+^ ST8SIA4-deficient neutrophils were comparable in both BM and blood (Figure S5G). These findings indicate that the monocytopenia in *St8sia4^-/-^*mice arises from the hematopoietic compartment and likely reflects a monocyte-intrinsic defect.

**Figure 4:**
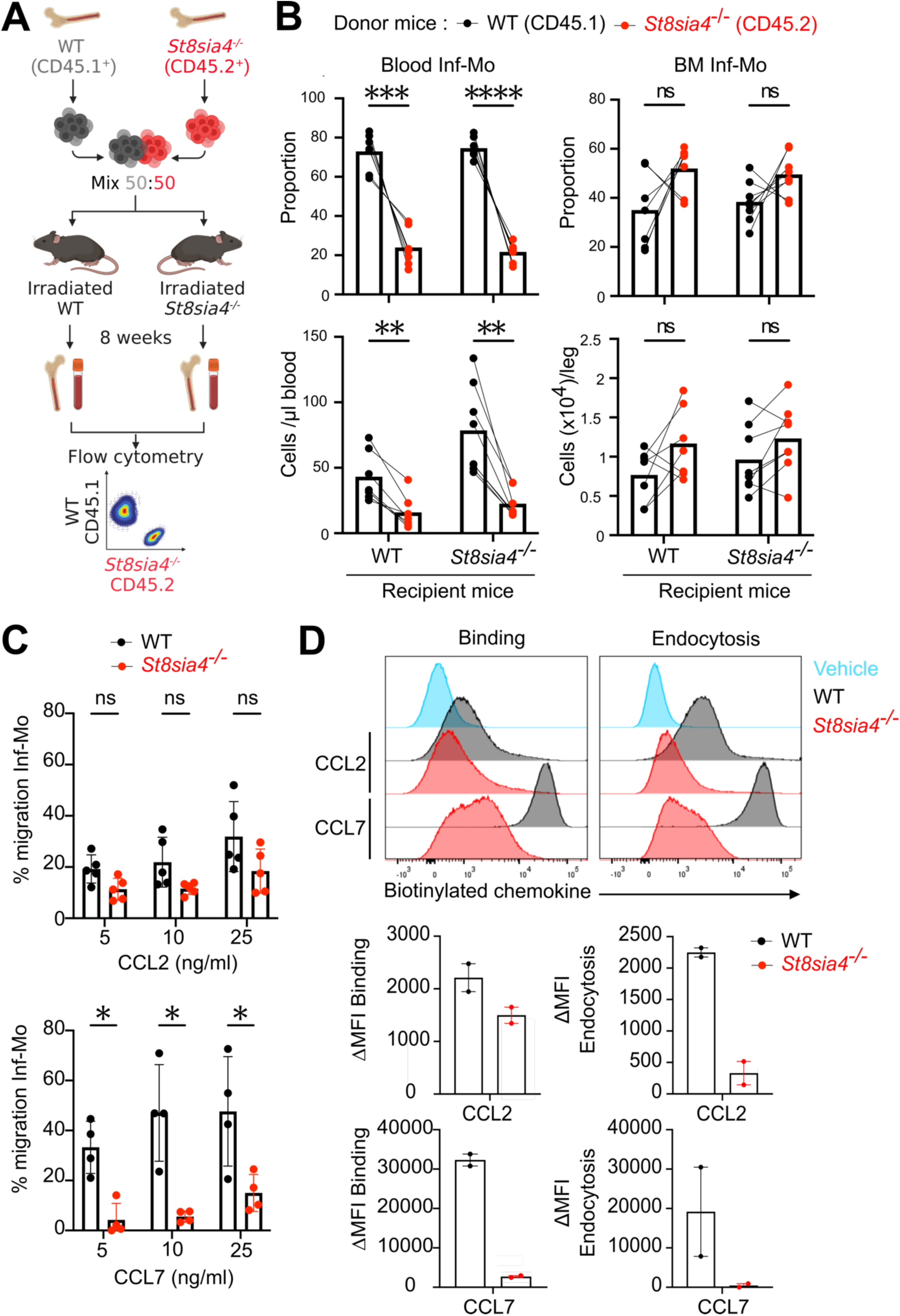
A monocyte-intrinsic defect in chemokine binding and endocytosis explains the monocytopenia in *St8sia4^-/-^* mice. (A) Schematic representation of mixed bone marrow chimera generation in mice. (B) Flow cytometry analysis of the proportion (top) and number (bottom) of inflammatory monocytes (Inf-Mo) derived from WT (CD45.1^+^) and *St8sia4^-/-^* (CD45.2^+^) donor mice in the blood (left panel) and BM (right panel) from irradiated WT and *St8sia4^-/-^*recipient mice. Each data point represents either donor CD45.1^+^ WT cells (black) or CD45.2^+^ ST8SIA4-deficient cells (red) (n=8 mice per group, pooled from 2 independent experiments). Connecting lines indicate that paired data originate from the same recipient mouse, while the bars represent the mean value across all mice Statistical analysis was performed using paired *t*-tests corrected for multiple comparison using Holm-Šídák method. **, *** and **** indicate adjusted *p*-values ≤0.01, ≤0.001 and ≤0.0001 respectively. The abbreviation ’ns’ stands for non-significant. (C) Percentage of inflammatory monocytes, purified from the bone marrow of WT and *St8sia4^-/-^* mice, that migrated *in vitro* in response to increasing concentrations of CCL2 or CCL7, as measured by a Transwell migration assay. Each data point corresponds to an individual experiment (n=4 independent experiments with pooled BM monocytes from 3-4 mice per group per experiment). Statistical analysis was performed using paired *t*-tests corrected for multiple comparison using Holm-Šídák method. * indicates a *p*-values ≤0.05. The abbreviation ’ns’ stands for non-significant. (D) Representative flow cytometry histograms (top) and bar plots (bottom) representing the delta geometric mean fluorescence intensity (ΔgMFI) of the binding and endocytosed of biotinylated CCL2 or CCL7, coupled to fluorescent streptavidin, to BM inflammatory monocytes isolated from WT and *St8sia4^-/-^* mice. ΔgMFI was calculated by subtracting the geometric mean of streptavidin group controls. Each data point corresponds to the ΔgMFI of an individual experiment (n=2, using pooled monocytes isolated from 3-4 mice per group). Bars and error indicate the mean ± SD.

To directly test this hypothesis, we isolated inflammatory monocytes from the BM of WT and *St8sia4^-/-^* animals and assessed their migration in response to CCL2 or CCL7 in transwell assays (Figures 4C and S5H). WT inflammatory monocytes responded robustly to increasing concentrations of either chemokine, with CCL7 inducing a stronger migratory response than CCL2. In contrast, migration in response to CCL7 was significantly reduced in ST8SIA4-deficient monocytes, whereas response to CCL2 was numerically, but not significantly, lower compared to WT cells (Figure 4C), consistent with our *in vivo* findings.

To investigate the basis of this impaired chemotaxis, we next assessed chemokine binding and endocytosis using biotinylated CCL2 and CCL7 complexed with fluorescently labeled streptavidin. Both CCL2 and CCL7 bound readily to WT monocytes (Figure 4D), whereas binding of CCL2 to ST8SIA4-deficient monocytes was slightly reduced, and CCL7 binding was nearly abolished. As a result, chemokine internalization was markedly reduced in ST8SIA4-deficient monocytes (Figure 4D).

Together, these findings suggest that monocytopenia in *St8sia4*^⁻/⁻^ mice is due to a monocyte-intrinsic defect in binding, internalizing, and responding to CCL2 and CCL7.

### NCAM1 is a major polySia carrier in inflammatory monocytes, and *Ncam1*^-/-^ mice also exhibit monocytopenia

Next, we aimed to characterize polysialylated proteins in inflammatory monocytes and to determine whether their absence could underlie the monocytopenia observed in *St8sia4*^⁻/⁻^ mice. Previous studies have identified several polysialylated proteins in immune cells, including NCAM1, NRP2, Golgi apparatus protein 1, and CCR7.^6,7,20,39^

Our analysis of polySia expression in BM inflammatory monocytes revealed two populations: one with high expression of polySia (polySia^high^) and another with lower expression (polySia^low^) (Figure 5A). Based on higher expression of CCR2 and lower expression of CXCR4 in the polySia^low^ population, these inflammatory monocytes appeared to be more mature than their polySia^high^ counterparts (Figure 5B).^37,40^

**Figure 5:**
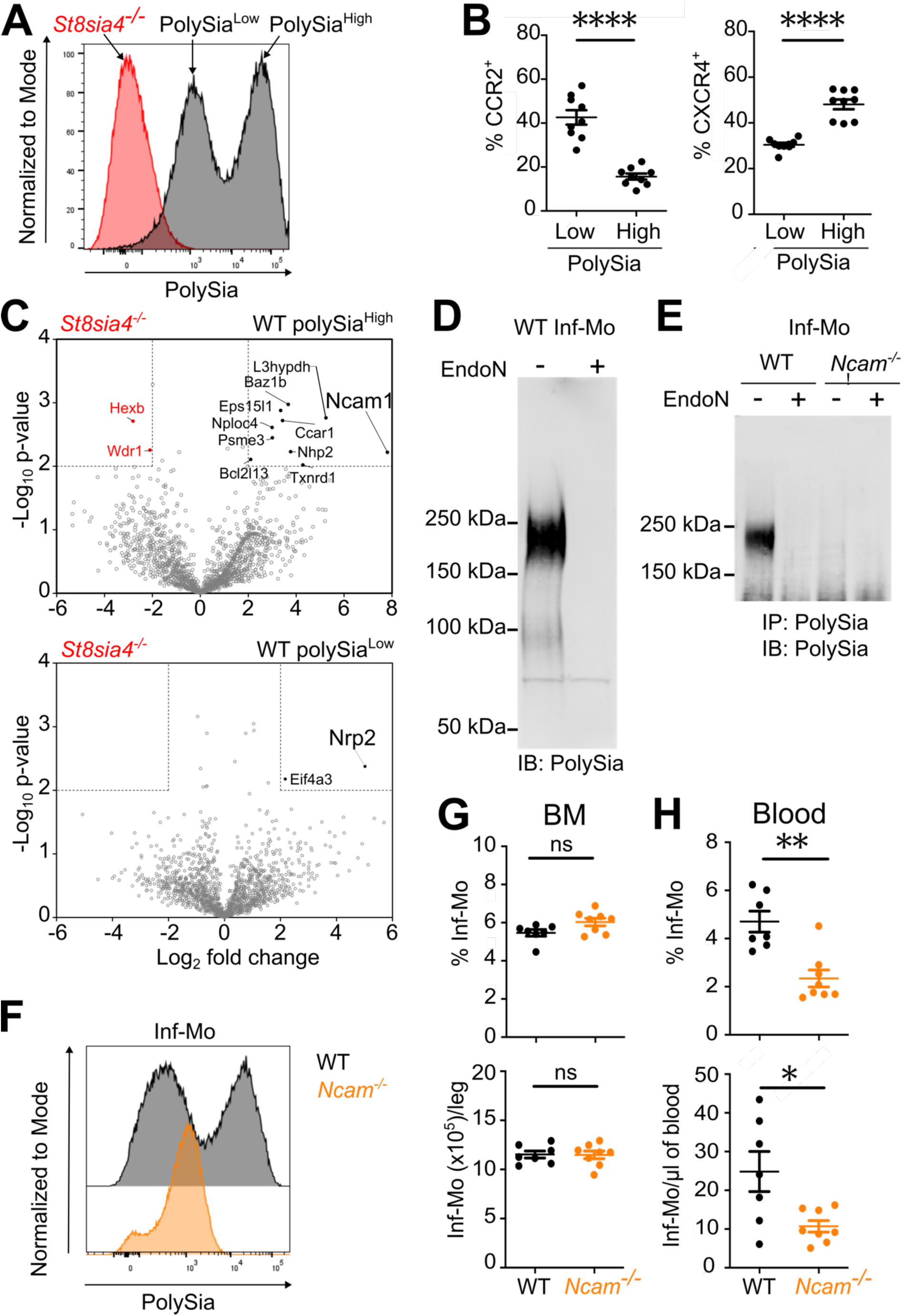
NCAM1 is a main polySia carrier protein in inflammatory monocytes and *NCAM^-/-^* mice also exhibit monocytopenia. (A) Representative flow cytometry histogram showing polySia surface expression on BM inflammatory monocytes from WT and *St8sia4^-/-^* mice. (B) Percentage of CCR2^+^ and CXCR4^+^ cells among BM inflammatory monocytes from WT and *St8sia4^-/-^* mice, as measured by flow cytometry. Each data point corresponds to an individual mouse (n=9 mice per group, pooled from 3 independent experiments). Bars and error indicate the mean ± standard error of the mean. Statistical analysis was performed using Welch’s *t*-tests. **** indicates a *p*-value ≤0.0001. (C) Label-free proteomic analyses of proteins immunoprecipitated using anti-polySia antibody from FACS-sorted polySia^high^, polySia^Low^ WT and *St8sia4^-/-^* BM inflammatory monocytes. Volcano plots showing the enriched proteins from polySia^high^ (top panel) or polySia^Low^ (bottom panel) inflammatory monocytes compared with *St8sia4^-/-^* cells. Statistical analysis of protein abundance values (log_2_ fold change, horizontal axis) was performed from different biological replicate experiments (n=3) using a paired sample *t*-test (-log_10_ *p*-value, vertical axis). Proteins found as significantly over or under-represented (*p* ≤0.01 and fold change ≥±4) are shown in black/red. (D) Western blot analysis of polysialylated proteins in WT and *St8sia4^-/-^*BM inflammatory monocytes showing the presence of a major band between 150 kDa and 250 kDa. (E) Western blot analysis of polysialylated proteins, treated or not with Endo-N-acetylneuraminidase (endoN), that were immunoprecipitated using anti-polySia antibody in WT and *Ncam1^-/-^* BM inflammatory monocytes. The presence of the major band between 150 kDa and 250 kDa in WT sample is not observed after endoN treatment or in *Ncam1^-/-^* samples demonstrating that this band corresponds to polysialylated NCAM1. (F) Representative flow cytometry histogram showing polySia surface expression on BM inflammatory monocytes from WT and *Ncam1^-/-^* mice. (G) Flow cytometry measurement of the frequency (top panel) and number of BM inflammatory monocytes (Inf-Mo) in WT and *Ncam1^-/-^* mice. Each data point corresponds to an individual mouse (n=7-8 mice per group). Bars and error indicate the mean ± standard error of the mean. Statistical analysis was performed using Mann-Whitney *t*-tests. The abbreviation ’ns’ stands for non-significant. (H) Similar to panel G for circulating inflammatory monocytes. Each data point corresponds to an individual mouse (n=7-8 mice per group). Bars and error indicate the mean ± standard error of the mean. Statistical analysis was performed using Mann-Whitney *t*-tests. * and ** indicate *p*-values ≤0.05 and ≤0.01 respectively.

To identify the polySia carrier proteins in these populations, we immunoprecipitated polysialylated proteins from FACS-sorted polySia^high^ and polySia^low^ inflammatory monocytes and analyzed the enriched proteins by proteomics. ST8SIA4-deficient inflammatory monocytes were used as negative controls. We detected a clear enrichment of NCAM1 in polySia^high^ and of NRP2 in polySia^low^ monocytes, both well-characterized polysialylated proteins (Figure 5C, and Table S2). Western blot analysis confirmed that NCAM1 is the main carrier of polySia in BM inflammatory monocytes (Figures 5D-E). The polySia^high^ monocyte population was completely absent in the BM of *Ncam1^-/-^* mice,^28^ although the total frequency and number of BM inflammatory monocytes were comparable to those in WT animals (Figures 5F–G). Similar to *St8sia4^-/-^* mice, *Ncam1^-/-^* mice displayed a marked reduction in circulating inflammatory monocytes, whereas eosinophil and neutrophil counts were normal (Figures 5H and S6A). These findings support an important role for NCAM1, the major polySia carrier in monocytes, in regulating their egress from the bone marrow.

### CCL2 and CCL7 do not directly bind to polySia

Previous studies demonstrated that polySia expressed on CCR7 acts as a coreceptor for the uptake of CCL21, but not CCL19, on DCs.^8,10,20,21^ Therefore, we hypothesized that polySia might be essential for the binding and uptake of CCL2 and CCL7, potentially explaining the chemotaxis defect of ST8SIA4-deficient monocytes. To test this, we performed *in vitro* chemokine binding assays in the presence or absence of soluble polySia as a binding competitor or endoneuraminidase to remove cell-surface polySia (Figure S6B). Both CCL2 and CCL7 bound to WT monocytes, but binding was not affected by the presence of soluble polySia or removal of polySia. As shown previously, binding of CCL7, and to a lesser extend CCL2, was decreased in ST8SIA4-deficient inflammatory monocytes. Similar to WT cells, binding was unaffected in the presence of soluble polySia or endoneuraminidase (Figures 6A-B). We confirmed the binding studies using Bio-layer Interferometry (BLI), in which we detected no interaction between CCL2 or CCL7 and polySia, although both chemokines interacted with heparan sulfate (Figures 6C and S6D). Binding of CCL21 to polySia and heparan sulfate were included as positive controls (Figure S6C). We also ruled out a role for GAGs in chemokine binding, as heparan and chondroitin sulfates were undetectable on the surface of BM inflammatory monocytes, and CCL7 binding remained unchanged following enzymatic digestion with heparinase or chondroitinase (Figures S6E-F).

**Figure 6:**
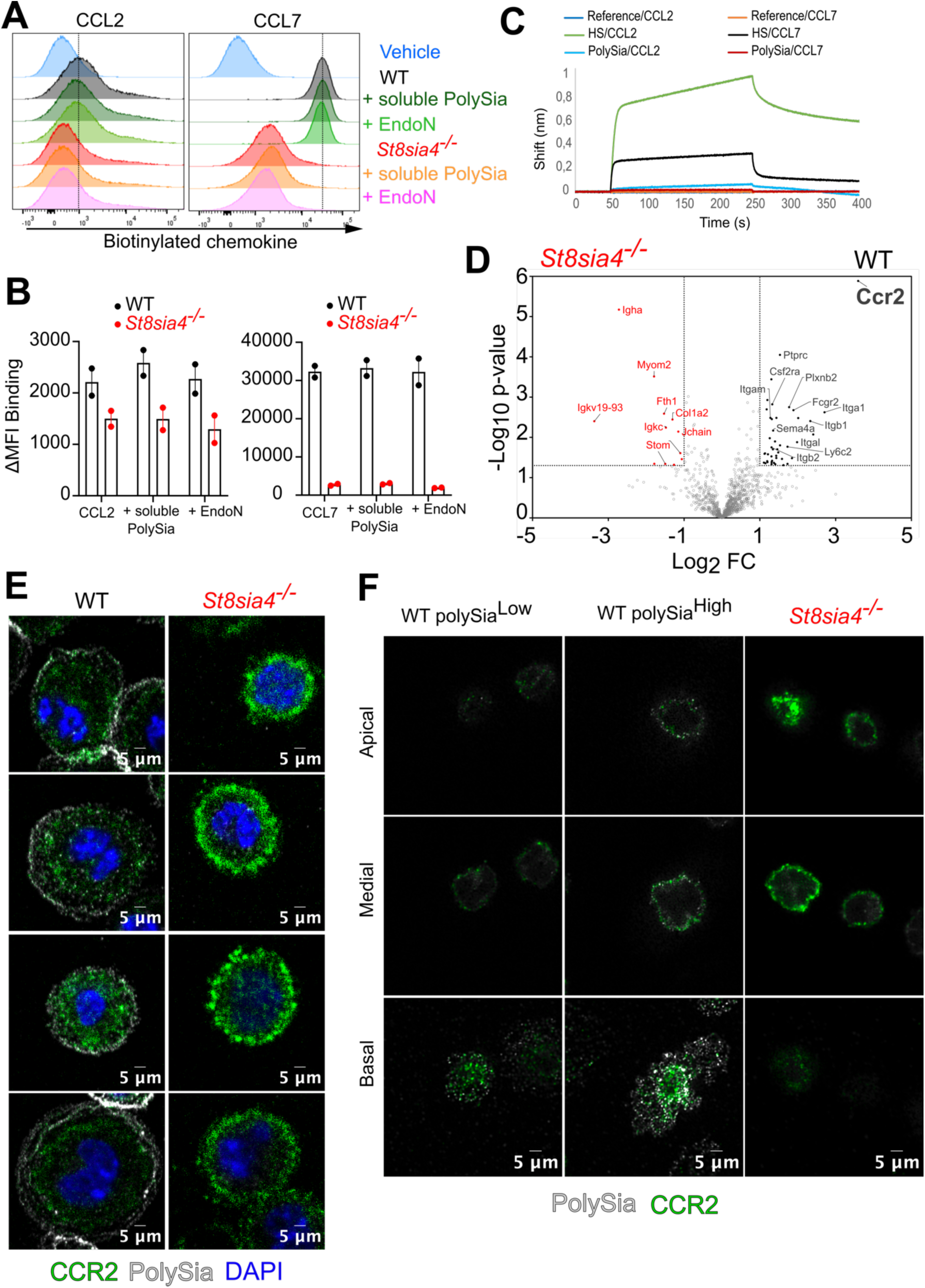
ST8SIA4-mediated polysialylation controls CCR2 surface localization and function but does not directly mediate CCL2/CCL7 binding. (A) Representative flow cytometry histograms of the binding of biotinylated CCL2 or CCL7, coupled to fluorescent streptavidin, to WT and *St8sia4^-/-^*BM inflammatory monocytes in the presence of an excess of soluble polySia (10 μg/ml) or after treatment of the cells with Endo-N-acetylneuraminidase (endoN) (n=2 independent experiments). (B) Bar plots representing the delta geometric mean fluorescence intensity (ΔgMFI) of the binding of biotinylated CCL2 or CCL7, coupled to fluorescent streptavidin, to WT and *St8sia4^-/-^*BM inflammatory monocytes in the presence of an excess of soluble polySia (10 μg/ml) or after treatment of the cells with endoN. ΔgMFI was calculated by subtracting the geometric mean of streptavidin group controls. Each data point corresponds to the ΔgMFI of an individual experiment (n=2, using pooled monocytes isolated from 3-4 mice per group). Bars and error indicate the mean ± SD. (C) Biolayer Interferometry (BLI) sensorgrams of mouse CCL2 and CCL7 (1 μM each) binding to polySia and heparan sulfate (HS). (D) Label-free proteomic analyses of proteins enriched using streptavidin coated beads after biotinylated CCL7-crosslinking on live WT and *St8sia4^-/-^* BM inflammatory monocytes. Volcano plots showing the enriched proteins from WT inflammatory monocytes compared with *St8sia4^-/-^* cells. Statistical analysis of protein abundance values (log_2_ fold change, horizontal axis) was performed from different biological replicate experiments (n=4) using a student *t*-test (-log_10_ *p*-value, vertical axis). Proteins found as significantly over or under-represented (*p* ≤0.05 and fold change ≥±2) are shown in black/red. (E) Images displaying cell surface CCR2 (green), polySia (white) and nucleus (DAPI in blue) in the basal region of WT and *St8sia4^-/-^* BM inflammatory monocytes, acquired using confocal microscopy. (F) Super-resolved images of cell surface CCR2 (green) and polySia (white) in low-adherent (non-spread) WT and *St8sia4^-/-^* BM inflammatory monocytes, acquired using random illumination microscopy. Optical sections (z-stack) corresponding to the basal, medial, and apical sections of the cells are shown.

These results show that, although ST8SIA4 and polysialylation regulate chemokine binding and internalization, polySia does not directly bind to CCL2 or CCL7.

### Loss of ST8SIA4 and polySia compromises CCR2 cell surface distribution and function

A majority of inflammatory monocytes in the BM and blood express CCR2 as the sole receptor for CCL2 and CCL7.^41^ Since polySia does not directly bind CCL2 and CCL7, and our earlier data showed that CCR2 expression is normal, or even increased, in ST8SIA4-deficient monocytes (Figure 3A, S7A), we hypothesized that the impairment of inflammatory monocyte mobilization might stem from a defect in CCR2 function.

To test this, we first confirmed that CCR2 is the primary receptor for biotinylated CCL7 on inflammatory monocytes using CCR2-deficient cells and flow cytometry (Figure S7B). Next, we performed a cross-linking experiment by incubating WT and ST8SIA4-deficient monocytes with an excess of biotinylated CCL7. Following verification of cross-linking efficiency by flow cytometry (Figure S7C), CCL7-protein complexes were enriched by affinity purification and analyzed by proteomics (Figure 6D and Table S3). We found that CCR2 was the most significantly enriched protein in WT monocytes after affinity purification of CCL7-protein complexes, whereas CCR2 was not detected in ST8SIA4-deficient cells under the same conditions (Figure 6D and Table S3). Our results thus confirm CCR2 as the main receptor of CCL7 and reveal that its function is compromised when ST8SIA4 and polySia are absent.

As CCR2 is present on the surface of ST8SIA4-deficient inflammatory monocytes but is unable to bind to CCL7, we examined its localization by confocal microscopy. Whereas WT inflammatory monocytes displayed uniform CCR2 staining at the basal region of the cell (Figure 6E), ST8SIA4-deficient monocytes exhibited more intense CCR2 staining, with receptor clusters arranged in circular patterns at the basal surface of the cell (Figure 6E). PolySia was also detected in the same regions as CCR2, but not colocalizing with it, in WT monocytes. Further analysis by super-resolution microscopy confirmed aberrant CCR2 distribution and receptor clustering in low-adherent ST8SIA4-deficient inflammatory monocytes (Figure 6F), with CCR2 distributed throughout the medial and apical regions of the cells and forming visible aggregates (Figure 6F). Of note, enzymatic removal of polySia from WT monocytes had little impact on cell surface distribution of CCR2, supporting the notion that cell surface polysialylation is not directly responsible for this phenotype (Figures S7D).

Collectively, these results indicate that ST8SIA4 deficiency alters the cell-surface distribution of CCR2, promotes receptor aggregation, and impairs its binding to CCL7.

As a consequence, inflammatory monocyte chemotaxis is markedly impaired, leading to defective egress of these cells from the bone marrow.

### ST8SIA4-Dependent Polysialylation Controls Cytoskeletal Organization in Monocytes

To further dissect the molecular mechanism by which ST8SIA4-dependent polysialylation supports CCR2 function, we performed a comparative proteomic analysis of inflammatory monocytes isolated from WT and *St8sia4^-/-^*mice (Figure 7A and tables S4). This analysis revealed significant differential expression (fold change ≥± 2; *p* ≤0.05) of multiple proteins related to the remodeling and polarization of the actin–myosin cytoskeleton in ST8SIA4-deficient monocytes, as confirmed by enrichment analysis (Figures 7A-B and tables S4), some of which were upregulated (Afdn, Myh4, Myl11, Acta1/Actc1, Ttn, Tnnt3, Neb, Sptan1, Arhgap45, Rhof, Tom1l2, Atp2a1, Rai14, Myl1, Dnm3) and others downregulated (Pak3, Sptb, Arhgap6, Srgap2, Prkg1, Tpm1, Pls1, Cald1, Nck2, Cavin2, Mylk, Lmna, Fhl1, Uxt, Plk1). Among the proteins upregulated in ST8SIA4-deficient monocytes were several involved in myofibril assembly (Myh4, Myl1, Myl11, Atp2a1, Acta1/Actc1, Neb, Ttn and Tnnt3; Figure 7A and table S4). We also noted differential expression of several proteins involved in cell adhesion (Cd302, Cdhr4, Cd226, Itgb6, Cd47) in ST8SIA4-deficient monocytes.

**Figure 7:**
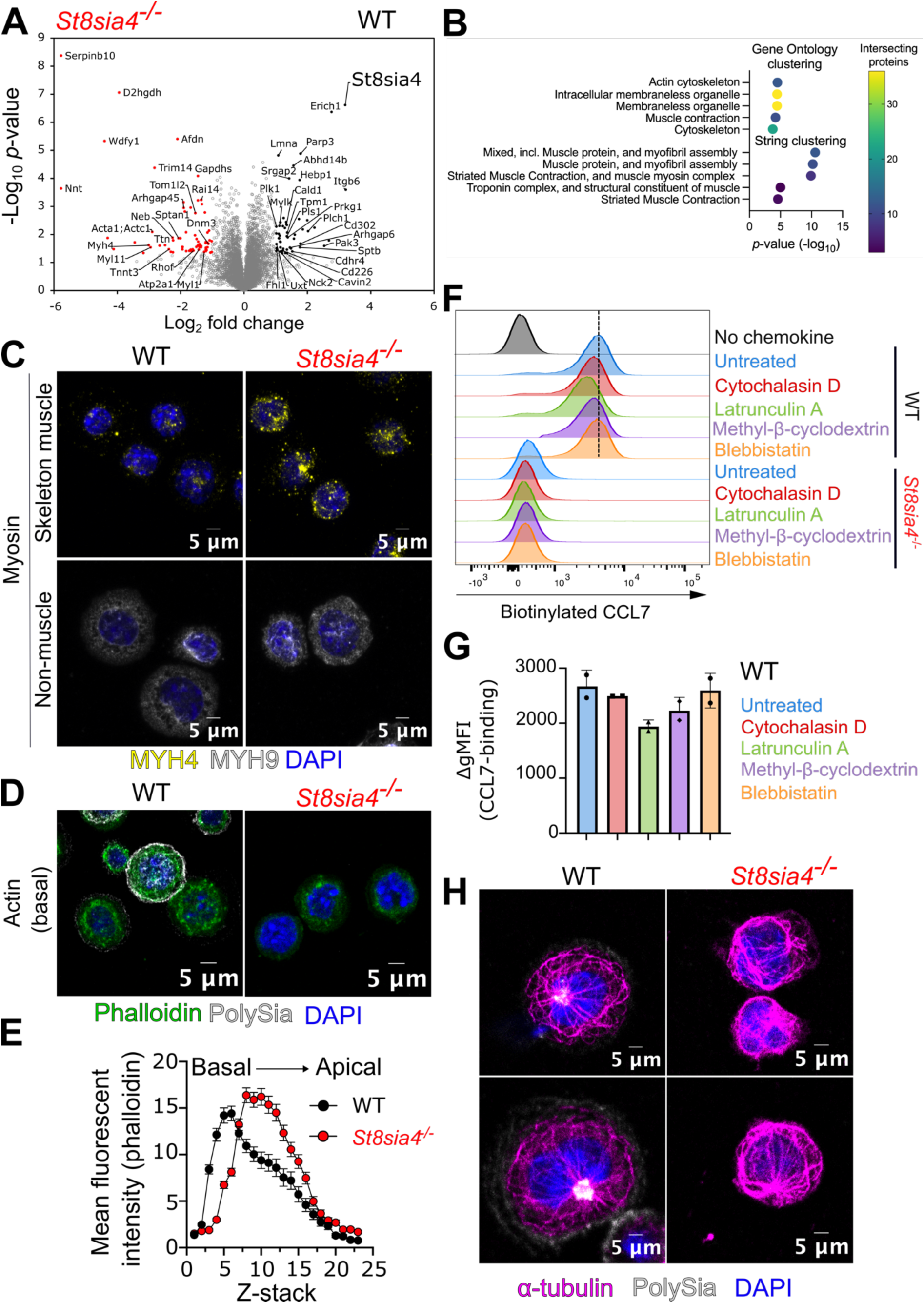
*St8sia4*^-/-^ inflammatory monocytes have important alteration of cytoskeletal architecture. (A) Label-free proteomic analyses of WT and *St8sia4^-/-^* BM inflammatory monocytes. Volcano plots showing the enriched proteins from WT inflammatory monocytes compared with *St8sia4^-/-^* cells. Statistical analysis of protein abundance values (log_2_ fold change, horizontal axis) was performed from different biological replicate experiments (n=4) using a student *t*-test (-log_10_ *p*-value, vertical axis). Proteins found as significantly over or under-represented (*p* ≤0.05 and fold change ≥±2) are shown in black/red. (B) Bubble plot showing the pathway enrichment analysis combining the proteins found significantly over or under-represented in *St8sia4^-/-^* BM inflammatory monocytes (Related to table S4). Gene Ontology (GO) and String cluster enrichment analysis was performed using g:Profiler and String app in Cytoskape. Only the top 5 GO terms among GO biological process, GO molecular function and GO cellular component categories are represented. Likewise, the top 5 String clusters are represented. The color of the dot represents the number of intersected proteins within a given term/cluster. (C) Confocal microscope images of skeleton muscle myosin (MYH4 in yellow), non-muscle myosin (MYH9 in white) and cell nucleus (DAPI in blue) in BM inflammatory monocytes from WT and *St8sia4^-/-^* mice. Images displaying MYH4 and MYH9 staining were generated using maximum-intensity projections. Images were processed using Fiji: ImageJ software. (D) Confocal microscope images of actin (phalloidin in green), polySia (in white), and cell nucleus (DAPI in blue) in WT and *St8sia4^-/-^* inflammatory monocytes. Images were processed using Fiji: ImageJ software and are representative of a single optical section corresponding to the basal region of the cells in contact with the lamella. (E) Quantification of the mean fluorescence intensity profile of actin (phalloidin staining) across optical sections (z-stack) representing the basal-to-apical region of WT and *St8sia4^-/-^* inflammatory monocytes. Mean fluorescent intensity quantification was performed on individual cells (n= 34-36 per group) over each optical section (Z-stack) using Fiji: ImageJ software. (F) Representative flow cytometry histograms of the binding of biotinylated CCL7, coupled to fluorescent streptavidin, to WT and *St8sia4^-/-^* BM inflammatory monocytes either untreated or treated with Cytochalasin D (1 μM; 15 minutes), Latrunculin A (1 μM; 15 minutes), Methyl-Δ-cyclodextrin (10 mM; 45 minutes), and (-)-Blebbistatin (20 mM; 45 minutes) (n=2 independent experiments with pooled monocytes isolated from 3-4 mice per group). (G) Bar plot representing the delta geometric mean fluorescence intensity (ΔgMFI) of the binding of biotinylated CCL7, coupled to fluorescent streptavidin, to WT and *St8sia4^-/-^* BM inflammatory monocytes either untreated or treated with Cytochalasin D (1 μM; 15 minutes), Latrunculin A (1 μM; 15 minutes), Methyl-Δ-cyclodextrin (10 mM; 45 minutes), and (-)-Blebbistatin (20 μM; 45 minutes). Each data point corresponds to an individual experiment (n=2, using pooled monocytes isolated from 3-4 mice per group). Bars and error indicate the mean ± standard deviation. (H) Confocal microscope images of microtubule network (a-tubulin in magenta), polySia (in white), and cell nucleus (DAPI in blue) in WT and *St8sia4^-/-^*inflammatory monocytes. Images were processed using Fiji: ImageJ software and are representative of a single optical section above the basal region of the cells.

ST8SIA4-deficient monocytes showed higher expression skeletal muscle myosin heavy chain IIb (MYH4) by confocal microscopy compared to WT cells (Figure 7C), whereas expression of non-muscle myosin heavy chain IIa (MYH9) was similar (by both proteomics and microscopy; Figure 7C, table S4). Analysis of the actin cytoskeleton using fluorescent phalloidin revealed a higher proportion of actin at the basal side of WT cells compared with ST8SIA4-deficient cells (Figures 7D-E). However, despite the alterations in the actomyosin cytoskeleton and differential expression of proteins involved in cell adhesion, adhesion of monocytes to various surfaces or extracellular matrix components was comparable in WT and ST8SIA4-deficient cells (Figure S7E).

Cytoskeletal remodeling and the organization of cholesterol-rich membrane microdomains critically regulate chemokine receptor conformation, nanoclustering, ligand-binding affinity, and ligand-dependent signaling.^42–46^ Because CCR2 was found to be mislocalized and clustered in ST8SIA4-deficient cells and the actin–myosin cytoskeleton was disrupted, we sought to evaluate the contribution of the cytoskeleton to CCR2–ligand interactions using pharmacological inhibitors, including cytochalasin D (blocks actin polymerization), latrunculin A (promotes actin depolymerization), methyl-β-cyclodextrin (depletes cholesterol, disrupts plasma membrane organization, and induces actin depolymerization), and (–)-blebbistatin (selectively inhibits myosin II). Of all drugs tested, only latrunculin A caused a modest reduction in CCL7 binding to WT cells by flow cytometry, although binding remained far higher than the low levels observed in ST8SIA4-deficient cells (Figure 7F). These results indicate that perturbation of the actin–myosin cytoskeleton or cholesterol-enriched membrane microdomains has little effect on CCL7 binding to CCR2 at the surface of WT monocytes. This suggests that the binding defect in ST8SIA4-deficient cells originates from upstream maturation steps that precede CCR2 delivery to the plasma membrane, including aberrant folding, altered post-translational modification or impaired intracellular trafficking.

The microtubule network plays a key role in organizing and regulating secretory pathways, directing protein transport from the endoplasmic reticulum through the Golgi apparatus to the plasma membrane.^47^ The microtubule cytoskeleton also regulates organization and positioning of intracellular organelles, in particular anchoring the Golgi apparatus near the microtubule-organizing center, thereby ensuring the proper assembly and maintenance of this organelle. Proteomics analysis revealed differential expression of proteins involved in microtubule assembly and intracellular protein folding (Cct6b, Cct8, Poc1a, Dnm3, Atxn7l1, Dhrsx, Sec11c) as well as in vesicle trafficking (Snca, Napg, Vamp3, Cavin2, Dnm3, Sft2d2, Mcfd2, Wdfy1) (Figure 7A and table S4). Accordingly, WT monocytes display more distinctly defined polarized microtubule bundles and meshworks than did ST8SIA4-deficient cells, as observed by confocal microscopy (Figures 7H and S7F). In addition, strong intracellular polySia labeling was detected in close proximity to the centrosome (Figures 7H and S7F). This is consistent with previous reports showing marked accumulation of polysialylated proteins, including NCAM1 and Golgi apparatus protein 1 (also known as E-selectin ligand 1), within the Golgi.^39^

Collectively, our results show that ST8SIA4-deficient monocytes display marked disruptions in actomyosin cytoskeletal organization and microtubule assembly. These defects likely underlie the observed CCR2 mislocalization, aggregation and dysfunction by impairing receptor processing and/or delivery to the plasma membrane.

## Discussion

The present study establishes polysialylation as a critical regulator of inflammatory monocyte chemotaxis and egress from the BM, with ST8SIA4-dependant polysialylation required for CCR2-dependent responses to cognate ligands. Loss of ST8SIA4-mediated polysialylation resulted in impaired egress of inflammatory monocytes from the BM, leading to monocytopenia both at steady state and during *Mtb* infection. The role of ST8SIA4 in controlling monocyte egress from the BM is likely conserved in humans, given that ST8SIA4 variants that promote upstream polyadenylation site usage (presumably diminishing full-length polysialyltransferase expression), are associated with lower circulating monocyte levels.

The degree of monocytopenia observed in *St8sia4^-/^*^-^ mice closely resembles that seen in *Ccl2^-/-^*or *Ccl7^-/-^* mice, although it does not reach the extent of blood monocyte reduction observed in *Ccr2^-/-^* mice.^35–37^ Notably, loss of ST8SIA4 did not fully abrogate egress of inflammatory monocytes from the BM, with *St8sia4^-/-^*cell accounting for approximately 20% of circulating inflammatory monocytes in mixed BM chimera experiments. The monocyte defect in *St8sia4^-/^*^-^ mice appeared to stem from a profound defect in CCR2-mediated chemotaxis, particularly in response to CCL7. The pronounced advantage of CCL7 over CCL2 in inducing monocyte chemotaxis in our models systems is interesting given that CCL7 has lower binding affinity for CCR2 and is only a partial agonist, resulting in reduced activation of G protein and β-arrestin signaling pathways, and decreased receptor internalization compared to CCL2.^48–50^ Of note, CCR2 has also been shown to function as a scavenger receptor for CCL2, promoting its clearance from the extracellular environment,^51–53^ an activity that appears to be independent of classical GPCR signaling components, such as G proteins, GPCR kinases, and β-arrestins. This scavenging activity might underly the differential responsiveness to CCL2 and CCL7 and perhaps also explain why the increase in serum CCL2 levels in *St8sia4^-/^*^-^ mice did not reach the magnitude observed in *Ccr2*^-/-^mice or following CCR2 antagonist treatment.^52,54^

One of the most well-documented functions of polySia is its positive role in cell motility and migration, as observed in different cell types and tissues.^55–60^ To date, two main mechanisms have been proposed to explain the role of polySia in cell migration. First, polySia, primarily carried by NCAM1, is proposed to act as a steric inhibitor of cell-cell and cell-extracellular matrix interactions, thereby facilitating cell detachment and migration. Second, polySia has been shown to directly interact with specific chemoattractant proteins (e.g., CCL21, FGF2, BDNF), enhancing chemotactic sensing. However, because ST8SIA4 deficiency did not increase inflammatory monocyte adhesion, and neither CCL2 nor CCL7 directly bound to polySia, our data suggest the existence of an additional mechanism by which polysialylation regulates monocyte chemotaxis and, more broadly, cellular migration.

Our findings reveal that ST8SIA4-deficient inflammatory monocytes exhibit pronounced disruptions in actomyosin cytoskeletal organization and microtubule networks. Notably, pharmacological perturbation of the actin–myosin cytoskeleton or cholesterol-rich membrane microdomains has minimal effect on CCL7 binding to CCR2 on WT inflammatory monocytes. Moreover, binding assays conducted at 4°C, which rapidly depolymerizes nearly all microtubules, further suggest that the defect observed in ST8SIA4-deficient cells likely arises from disturbances in early cellular processes preceding plasma membrane localization, such as receptor folding, post-translational modification (e.g., glycosylation, tyrosine-sulfation), and/or exocytosis. Supporting this hypothesis, we detected intracellular polysialylation concentrated near the microtubule-organizing center, consistent with previous reports of abundant polysialylated proteins in the adjacent Golgi apparatus.^39^ Because NCAM1 contributes to the Golgi-resident polySia pool, and because *Ncam1^-/^*^-^ mice phenocopy the monocytopenia seen in *St8sia4^-/^*^-^ mice, these findings collectively implicate polysialylated NCAM1 as a regulator of actomyosin and microtubule organization and/or of CCR2 function and trafficking to the plasma membrane in inflammatory monocytes. Consistent with this model, NCAM1 engages multiple cytoskeletal elements—including α- and β-tubulin, α-actinin-1, tropomyosin and β1-spectrin^61–63^, several of which were differentially expressed in ST8SIA4-deficient inflammatory monocytes relative to WT cells (Table S4). NCAM1 also interacts with Kinesin-1, which drives its microtubule-dependent trafficking from the *trans*-Golgi network to targeted regions of the plasma membrane.^64,65^ In neurons, NCAM1 further stabilizes microtubule projections at synapses and promotes the delivery of *trans*-Golgi network delivery of synaptic proteins to the cell surface.^66,67^ Together, these findings point to a mechanistic link between ST8SIA4-dependent polysialylation, NCAM1 and CCR2 localization and function in inflammatory monocytes, a relationship that warrants further investigation.

In addition to inflammatory monocytes, ST8SIA4-deficient hematopoietic progenitors have also been reported to exhibit defects in BM egress.^22^ As it is now well established that hematopoietic progenitors migrate in a CCR2-dependent manner,^68^ our findings may help to explain the impaired BM egress of these cells in *St8sia4^-/-^* mice. Future work will be required to confirm this hypothesis, and more generally, to assess whether this dysfunction extends to other chemokine receptors across diverse cell types.

The human data support a conserved role for ST8SIA4 in regulating monocyte release into the blood stream and highlight a potential role for alternative isoform usage in the regulation of ST8SIA4 availability. Our observations suggest that multiple genetic variants may independently alter promoter choice, splicing and polyA usage, with a combined impact on the availability of the full-length, canonical isoform of the enzyme. Given that premature termination of transcription would lead to an incomplete glycosyltransferase domain, we expect the observed change in isoform usage to decrease total ST8SIA4 activity, which could explain the reduced circulating monocyte counts. Further characterization of the full isoform repertoire of *ST8SIA4* by long read sequencing and systematic exploration of haplotypic effects on isoform usage will be required to understand the genetic basis of ST8SIA4 expression in humans.

The genetic data also suggest an effect of ST8SIA4 on lymphocyte counts, and highlight the functional relevance of ST8SIA4 in the etiology of lupus, possibly related to ST8SIA4’s ability to regulate migration of classical monocytes to the sites of inflammation.^69^ Further study is warranted to investigate the role of ST8SIA4-dependent polysialylation in these phenotypes and to dissect the underlying mechanisms.

In summary, our study establishes polysialylation as a key regulator of inflammatory monocyte chemotaxis and CCR2 function, uncovering a previously unrecognized layer of control in immune cell trafficking. By demonstrating that ST8SIA4-dependent polysialylation regulates monocyte egress from the bone marrow, both at steady state and during infection, we reveal a fundamental mechanism that bridges glycosylation, cytoskeletal dynamics, and chemokine receptor function. These findings not only advance our understanding of monocyte biology but also open new avenues for targeting monocyte mobilization in inflammatory diseases, infections, and autoimmune disorders like systemic lupus erythematosus. Future work exploring the broader role of polysialylation in other immune cell types and its therapeutic potential may reveal new avenues to modulate immune responses in health and disease.

### Limitations of the study

Although our findings firmly establish ST8SIA4-dependent polysialylation is critical for CCR2-mediated monocyte egress, several points warrant further exploration. First, our mechanistic studies rely primarily on well-validated mouse models, and complementary work in human systems will help extend these insights to human biology. Second, while we confirm NCAM1 as the dominant polySia carrier in inflammatory monocytes, additional polysialylated proteins, most notably NRP2, may operate in parallel or in specific contexts, a possibility that does not diminish the central role identified here. Third, the cytoskeletal and receptor-organization changes observed in the absence of polysialylation highlight a clear mechanistic link, even as the full molecular details remain to be refined. Finally, although human genetic associations at the *ST8SIA4* locus point to potential relevance in diseases such as SLE, establishing causality and therapeutic implications will require dedicated future studies. Together, these considerations represent natural extensions of our work rather than constraints on the conclusions presented here.

## Supplemental information

Supplemental figures and tables can be found in separate files.

## Acknowledgements

We thank Céline Beronne, Flavie Moreau, Thomas Herpin, and Aline Tridon (Genotoul Anexplo-IPBS platform, Toulouse, France) for mouse care. We also extend our gratitude to several IPBS members including Emmanuelle Näser and Eve Pitot for assistance with flow cytometry and confocal microscopy, Renaud Poincloux for his scientific advices, Stella Rousset for helping purifying monocytes, Benjamin Raymond for helping with biochemistry, Olivier Saurel for NMR analysis of biotinylated polySia and Arnaud Métais for assistance with epifluorescence microscopy. We gratefully acknowledge Julie Noguerol and Olivier Joffre (Infinity, Toulouse, France) for their invaluable assistance with irradiation procedures for mouse chimera experiments. We thank the Genotoul TRI-IPBS imaging facility, member of the national infrastructure France-BioImaging supported by the French National Research Agency (ANR-10-INBS-04; ANR-24-INBS-0005 FBI BIOGEN). This project was financially supported by the Centre National de la Recherche Scientifique, Université de Toulouse, the Agence Nationale de la Recherche (grants ANR-17-CE11-0006 and ANR-23-CE44-0001 to Y.R., and grant ANR-22-CE12-0030-01 to M.R.), Fondation pour la Recherche Médicale (to A.B.) and the Fondation Bettencourt Schueller (grant Explore-TB to O.N.). This work also used the platforms of the Grenoble Instruct Centre (ISBG; UMS 3518 CNRS-CEA-UJF-EMBL) with support from French Infrastructure for Integrated Structural Biology (ANR-10-INSB-05-02) and Grenoble Alliance for Integrated Structural Cell Biology (ANR-10-LABX-49-01) within the Grenoble Partnership for Structural Biology. H.H. received support from the Deutsche Forschungsgemeinschaft (DFG, German Research Foundation), project numbers 324633948 (grant Hi 678/9-3) and 409784463 (grant Hi 678/10-2, FOR2953).

## Authorship Contributions

Conceptualization: A.B., O.N. and Y.R.; Data curation: A.B., N.G., R.S., A.GdeP, M.R. and Y.R.; Formal analysis: : A.B., N.G., R.S. and Y.R. (mouse) and M.R and L.Q-M. (Human); Funding acquisition: Y.R.; Investigation: A.B, N.G., D.H., S.M., Y.R. (mouse experiments), M.R and L.Q-M (Human genetic), A.B. and Y.R. (Biochemistry), Y.R., A.S., K.C., O.B-S and A.GdeP. (Proteomics), R.S. and H.L-J (BLI). Methodology: A.B., N.G., D.H., O.N., and Y.R.; Project administration: O.N., and Y.R.; Resources: R.S. and H.L-J (recombinant heparinases), A.M-K and H.H. (*St8sia4*^-/-^ and *Ncam1*^-/-^ mice; recombinant endoneuraminidase), N.A. (*Ccr2*^-/-^ mice). Supervision: Y.R.; Validation: A.B., N.G., M.R., R.S. and Y.R.; Visualization: A.B., N.G., M.R., C.G., R.S. and Y.R.; Writing – original draft: Y.R.; Writing – review & editing: A.B., M.R., A.GdeP., H.H., R.S., H.L-J., O.N., and Y.R.

## Declaration of generative AI and AI-assisted technologies in the manuscript preparation process

During the preparation of this work the author(s) used ChatGPT (OpenAI) in order to improve language and readability. After using this tool/service, the author(s) reviewed and edited the content as needed and take(s) full responsibility for the content of the published article.

## Methods

### Mice

Animal care and experimentation consistent with the French Policy for Animal Protection and were approved by the Ethic Committee of Toulouse Biological research Federation (C2EA-01) and the Ministry of Higher Education and Research (Agreement APAFiS #26896). In particular, all efforts were made to minimize animals’ discomfort and suffering. *St8sia4*^-/-^ and *Ncam1^-/-^* mice were kindly provided by Prof. H. Hildebrandt (Hannover Medical School, Germany)^18^ . C57BL/6J and *St8sia4*^-/-^ mice were housed and bred in a specific pathogen-free environment at the animal facility of the Institut de Pharmacologie et de Biologie Structurale, Toulouse, France. *Ncam1^-/-^*were housed and bred at the central animal facility of Hannover Medical School. *CCR2^-/-^* mice were housed and bred at Universidad de Zaragoza.

### Mouse infection and lung preparation

*Mycobacterium tuberculosis* (*Mtb*) H37Rv strain was grown in Middlebrook 7H9 medium (Difco) supplemented with Middlebrook ADC (BD #211887) and 0.05% Tween® 80 (Sigma-Aldrich #P4780). For infection, growing *Mtb* at exponential phase was centrifuged at 3000 x g and resuspended in Dulbecco’s phosphate-buffered saline, no calcium, no magnesium (DPBS, Gibco^TM^, ThermoFisher scientific #14190144). Bacterial aggregates were dissociated by twenty passages through a 26G blunt needle. Bacterial suspension concentration was determined by measuring the optical density at 600 nm, and then resuspended in DPBS for *in vivo* infection. Female C57BL/6J or *St8sia4*^-/-^ mice (6- to 12-week-old) were anesthetized by an intraperitoneal injection of ketamine (Virbac) and xylasine (Bayer). Mice were infected intranasally with 1000 bacilli of *Mtb* diluted in 25 μL of DPBS. The inoculum bacterial load (colony forming units) was confirmed by plating different dilutions on Middlebrook 7H11 Agar Base (Sigma-Aldrich #M0428) supplemented with Middlebrook OADC (BD #211886). The plates were incubated at 37°C for 3 weeks before enumeration of colonies. At 7-, 14-, 21- or 42-days post-infection, mice were anesthetized in gas chambers containing isoflurane and received a retro-orbital injection of 100 μL of DPBS containing 2 μg of Alexa Fluor® 700 anti-mouse CD45.2 (BD Biosciences # 560693) to discriminate between parenchymal and intravascular cells in subsequent flow cytometry analyses. After 5 min, mice were euthanized by cervical dislocation followed by dissection to harvest the lungs. Lungs were homogenized in HBSS, no calcium, no magnesium, no phenol red (HBSS, Gibco^TM^, ThermoFisher scientific #14175095) using a gentleMACS dissociator (C Tubes, Miltenyi), and then incubated with DNAse I (0.1 mg/mL, Roche) and collagenase D (2 mg/mL, Roche) for 30 min at 37°C under 5% CO2. Pulmonary bacterial loads were measured by plating serial dilutions of the lung homogenates onto Middlebrook 7H11 OADC selective agar. Plates were incubated at 37 °C and colonies were enumerated three weeks after. The remaining lung homogenate fraction was filtered using a 40 μm cell strainer (Dutscher #141378C) on a sterile 50 mL conical tube, and centrifuged at 1 400 rpm for 5 min. Pelleted cells were resuspended in a red blood cell (RBC) lysis buffer (150 mM NH_4_Cl, 10 mM KHCO3, 0.1 mM EDTA pH 7.2), centrifuged at 1 400 rpm for 5 min, and then processed for flow cytometry analysis (see below).

### Isolation of bone marrow, spleen and peripheral blood leukocytes

To prepare bone marrow (BM) cells, mice were euthanized by cervical dislocation under isoflurane anesthesia and bones from the two hind legs (hips, femurs and tibiae) were harvested. Bones were soaked in 70% ethanol and then incubated for few minutes in DPBS. Bones were cut at both ends using sharp scissors and placed into a sterile 500 µL microtube with a pre-punched hole. Tubes containing the bones were placed into a 1.5 mL microtube and the double-layered tubes were shortly centrifuged using a microcentrifuge in order to pellet the bone marrow. The latter was resuspended in 1 mL of DPBS supplemented with 0.5 % fetal bovin serum (FBS, Sigma-Aldrich #F7524) and 2 mM EDTA (Euromedex #EU0084-B) followed by filtration on a 40 μm cell strainer. The rest of the bones was crushed with scissors in 2 ml of DPBS/ 0.5 % FBS/ 2 mM EDTA. The suspension was filtering on a 40 μm cell strainer and combined with the bone marrow. Red blood cells were lysed by addition of RBC lysis buffer for 5 minutes and the BM suspension was centrifuged at 1 400 rpm for 5 minutes. Pelleted cells were either directly stained with antibodies for flow cytometry analysis (see below) or used for the isolation of BM inflammatory monocytes. The latter were purified by negative selection using a monocyte isolation kit (Miltenyi Biotec #130-100-629) according to the manufacturer’s instructions, taking into account 1.10^9^ cells per BM suspension.

Regarding splenocyte isolation, mice were anesthetized by an intraperitoneal injection of ketamine (Virbac)/xylasine (Bayer), subjected to a cardiac perfusion with 20 mL of DPBS using 25G needle, and then euthanized by cervical dislocation. Spleen was then harvested, placed onto a 70 µm cell strainer (Dutscher #141379C) on a sterile 50 mL conical tube, and crushed using the flat end of a syringe plunger. The cell strainer was then rinsed with at least 10 mL of DPBS to release the splenocytes.

In order to isolate peripheral blood leukocytes, mice were anesthetized by an intraperitoneal injection of ketamine (Virbac)/xylasine (Bayer). Blood was collected by cardiac puncture using 25G needle and a 1 mL syringe and immediately transferred into a microtube containing 0.5 mM EDTA (10% of expected blood volume) to prevent coagulation. Mice were then euthanized by cervical dislocation. Red blood cells were lysed by addition of RBC lysis buffer for 5 min. Leukocytes were collected by centrifugation at 1 400 rpm for 5 min, and further processed for flow cytometry analysis.

### Flow cytometry and data acquisition

Between 5 x 10^5^ to 1 x 10^6^ cells from organs or *in vitro* cultures were transferred in a V-bottom 96-well culture plate. The plate was centrifuged at 600 x g for 3 minutes, and the supernatant was removed by flicking. Cells were then resuspended in 30 μL of antibody cocktail, diluted in cell staining buffer (BioLegend; #420201) or FACS buffer (DPBS/ 2% FBS/ 2mM EDTA), and incubated for 30 minutes at 4 °C. Following staining, cells were washed with 170 μL FACS buffer and finally resuspended in 200 μL FACS buffer. All these steps are interspersed with pelleting of the cells by centrifuging the plate at 600 x g for 3 minutes at 4°C, followed by removal of the supernatant by flicking the plate. Cells were transferred to round-bottom tubes, diluted to a final volume of 600 µL, and analyzed by flow cytometry. In all cases the antibody cocktail was composed of TruStain FcX™ antibody solution (1:50 dilution; Biolegend; # 101320) to inhibit non-specific antibody binding to FCyR, and Live/Dead (Invitrogen; #L23105) to exclude dead cells. Additional antibodies used for flow cytometry analysis are listed in table S4. Intracellular cell staining was performed after the staining of cell surface marker described above. Briefly, cells were permeabilized and fixed using the Foxp3/Transcription Factor Staining Buffer Set kit (Invitrogen; #00-5523-00). Following fixation (using the fixation solution provided in the kit) for 30 min at room temperature, cells were incubated at room temperature for 45 minutes in 30 μl antibody cocktail diluted in permeabilization buffer (provided in the kit), washed with 170 μl FACS buffer (DPBS/ 2% FBS / 2mM EDTA), resuspended in 200μl FACS buffer and transferred to round-bottom tubes for the flow cytometry analysis. All these steps are interspersed with pelleting of the cells by centrifuging the plate at 700 × g for 3 minutes, followed by removal of the supernatant by flicking the plate Cells obtained from tissues infected by *Mtb* were stained with antibody as described above and then fixed with a 4% paraformaldehyde solution for 2 hours at room temperature. Active caspases-3/-7 are detected using the Vybrant Caspase-3 and -7 assay kit (FITC; Molecular Probes; #V35118) following the supplier’s instructions, then labelled with the appropriate antibodies as described above. CountBright absolute counting beads (Life technologies #C36950) are used for absolute cell counting by flow cytometry, following the supplier’s instructions. Cells were acquired on a Fortessa X20 flow cytometer (BD Biosciences) and data were analyzed using Flowjo v10 data analysis software.

### Analysis of the concentration chemokines and growth factor

To obtain serum, mouse blood was collected by cardiac puncture as described above. The blood was then left at room temperature for 30 minutes (without EDTA) followed by centrifugation at 1000 x g for 20 minutes. The serum (corresponding to the supernatant) was collected. For the recovery of BM supernatants, two long bones from the hind legs (1 femur and 1 tibia) were recovered from each mouse, as previously described. Following centrifugation of BM suspension for 5 minutes at 400 x g at 4°C, the BM supernatant was collected. The various factors involved in myelopoiesis were then quantified using the LEGENDplex Mouse HSC Myeloid panel kit (#740683; Biolegend). The LEGENDplex was then performed and analyzed according to the supplier’s instructions. Measurement of serum concentration of CCL2 and CCL7 in WT and *St8sia4*^-/-^ mice was performed using the MCP-1 Mouse ELISA Kit (Invitrogen #BMS6005) and MCP-3 mouse Instant ELISA Kit (Invitrogen #BMS6006INST; Invitrogen) according to the supplier’s recommendations.

### Mixed bone marrow chimeras

WT CD45.2 and *St8sia4*^-/-^ CD45.2 recipient mice (6-week-old) were irradiated using a split irradiation dose (2 x 550 rad). The following day, bone marrow from WT CD45.1 and *St8sia4*^-/-^ CD45.2 mice (from 8 to 12 weeks old) were isolated as indicated above. An equal number of WT CD45.1 and *St8sia4*^-/-^ CD45.2 was mixed together (2.5 x 10^6^ cells of each) in DPBS and 150µl with 5 x 10^6^ cells were injected intra-orbitally into recipient WT CD45.2 or *St8sia4*^-/-^ CD45.2 mice. Mice were treated with 300 µl of Adjusol® in 100 ml drinking water every day for at least 3 weeks. After 8 weeks, mice were sacrificed, peripheral blood and BM were harvested and quantitated by flow cytometry as described above.

### *In vivo* and *in vitro* chemotaxis assays

To analyze the BM egress of monocytes *in vivo*, we performed a retro orbital injection of 100 µL of DPBS alone (vehicle) or DPBS containing CCL2 (500 ng) or CCL7 (250 ng) in WT and *St8sia4*^-/-^ animals. After 30 min, mice were euthanized followed by isolation and flow cytometry analysis of blood and BM leukocytes as described previously.

In order to measure the chemotaxis of WT and *St8sia4*^-/-^ BM monocytes *in vitro*, we used a transwell migration assay. In brief, 2 x 10^5^ monocytes in 100 μl of medium without FBS (RPMI GlutaMAX medium / 1% Penicillin Streptomycin (PS)) were placed onto the membrane of the 24-well transwell inserts (8 µm pore size, Falcon; #353097) and incubated for 1 hour at 37°C to allow them to settle. Then, 600 μl of migration buffer (RPMI 1640 GlutaMAX medium / 1% PS / 10% FBS) containing increasing concentration (0, 5, 10, 25 ng/ml) of CCL2 (Peprotech; #250-10,) and CCL7 (Peprotech; #250-08) were added to the well directly below the transwell insert. After incubation for 2 hours at 37°C, transwells were removed and cells that have migrated into the well were allow to adhere for 1 hours at 37°C. As per the manufacturer’s instructions, cells that migrated into the lower chamber were lysed with CellTiter-Glo 2.0 Reagent (#G9241; Promega) followed by luminescence measurement on a CLARIOstar. All assays were performed in quadruplicate. Results were expressed as the percentage of cells that migrated relative to the initial inoculum, after subtracting basal migration toward the medium without chemokine.

### Measurement of chemokine binding and endocytosis on cells

Monocytes were enriched from the BM of WT or *St8sia4*^-/-^ mice as described above and 5 x 10^5^ monocytes in 200 μL RPMI containing 10 % FBS were transferred in a V-bottom 96-well plate. Cells were incubated for 30 min with either streptavidin-PE (#405204, BioLegend) (negative control) or with 20 nM of CCL2-biotin (#ABIN6952158, antibodies online) or CCL7-biotin (#ABIN6952208, antibodies online) pre-conjugated with streptavidin-PE (molar ratio 1:4). Incubation was performed either at 4°C or at 37°C in 5% CO_2_ incubator to analyse chemokine binding or endocytosis respectively. Cells were then pelleted by centrifugation at 600 x g for 3 minutes, and washed once with 200 μL of Flow cytometry buffer (DPBS supplemented with 0.5 % FBS and 2 mM EDTA). Following another centrifugation step, cells were resuspended in 200 μL of Flow cytometry buffer, and finally transferred to round-bottom tubes. Flow cytometry analysis was performed either on the same day without cell fixation or the following day after fixing the cells with 4% paraformaldehyde. Glycosidase digestion was performed, prior to chemokine staining, by incubating the cells in V-bottom 96-well plate with either 1 µg/mL of endoneuraminidase,^70^ 0.5 U/mL of Chondroïtinase ABC (#C3667, Sigma-Aldrich), or a mix of Heparinase I at 0.1 U/mL and Heparinase III at 0.1 U/mL for 1h at 37°C under gentle agitation. Efficacy of digestion was confirmed by flow cytometry either on monocyte (for polySia; Figure S6B) or on Mouse Embryonic Fibroblasts (for heparinases). Competition using soluble PolySia was done by adding 10 µg/mL of soluble polySia (Interchim, #628897) to the chemokine/streptavidin-PE mix. For drug treatment, cells were incubated in presence of 1 μM Cytochalasin D (#5.04776, Sigma-Aldrich) or 1 μM Latrunculin A (#428021, Sigma-Aldrich) for 15 minutes, or 20 μM (-)-Blebbistatin (#203391, Sigma-Aldrich) or 10 mM methyl-β-cyclodextrin (#C4555, Sigma-Aldrich) Latrunculin A (#428021, Sigma-Aldrich) for 45 minutes at 37°C in CO2 incubator. Cells were washed twice with in 200 μL of DPBS to remove residual enzymes, glycan fragments or drugs, and then incubated with the chemokine/streptavidin complexes as described previously.

### Progenitors CFU

For GMP purification, the BM cells were isolated from the 2 hind legs (femur, tibia and hip) as explained previously. Hematopoietic cells were then enriched by negative selection using the mouse lineage cell depletion kit (Miltenyi; # 130-110-470) according to supplier’s instructions. The remaining cells were then resuspended in Cell Staining Buffer (CSB, BioLegend; #420201) and incubated for 30 minutes at 4°C with an antibody cocktail containing FITC-coupled antibody targeting CD3, CD19, B220, Ter-119, CD49b, Ly6G, CD11c, and CD11b to exclude lineage cells and additional fluorescent antibodies targeting CD117 (C-kit), CD16/32 (FcyR), Ly-6A/E (Sca-1), FLT3, CD34, CD115, Ly6C for GMP gating (see figure S3A). GMP sorting was performed by flow cytometry using an Aria Fusion (BD Biosciences) and cells were collected in 15 mL canonical tubes pre-coated with FBS. After sorting, cells were diluted with 2 mL Iscove’s MDM with 2% FBS (StemCell technologies; # 07700), pelleted by centrifugation at 300 x g for 10 minutes and resuspended in 1 mL Iscove’s MDM with 2% FBS. Following sorting, 0.25 x 10^3^ GMPs were seeded in triplicate in 6-well plate Methylcellulose-based medium with recombinant cytokines (Stem Cell technologies; # 03534) containing 1% PS to allow the generation of CFU-M (Colony Forming Unit-macrophages), CFU-G (-granulocytes) and CFU-GM (mix of - macrophages and granulocytes) in 6-well plate. After 10-14 days of culture, the wells were scanned using an EVOS M7000 microscope (Invitrogen) and colonies were identified and counted in each well blindly by 2 independent individuals. Colonies are identified based on morphological criteria as described by (Olsson et al., 2016). Briefly, CFU-M form diffuse colonies composed of large cells with an oval/round shape; CFU-G are small dense colonies composed of small cells (compared to CFU-M); CFU-GM form large colonies with generally several dense centers and are composed of granulocytes and macrophages.

### Biolayer Interferometry (BLI) assays of chemokines binding on PolySia and GAG (HS)

BLI studies were performed using a FortéBio OctetRED96e instrument from Sartorius and collected data were recorded with the manufacturer’s software (Data Acquisition HT 11.0). Chemokines (200 nM of CCL21, 1 µM of CCL2 or CCL7), PolySia and HS were resuspended in 0.01 M HEPES pH 7.4, 0.15 M NaCl, 0.005% v/v Surfactant P20 (HBS-P+) buffer (Cytiva). Samples (200 μL) were deposited in wells of 96-well plates (Nunc F96 Microwell, ThermoFisher Scientific), at 25 °C, under agitation (1000 rpm) during all the run. All biosensors were pre-wetted in 200 μL of HBS-P+ buffer for 10 min before each test. For the capture, SAv-coated biosensors (Sartorius) were loaded with biotinylated PolySia or biotinylated HS for 600 s. The concentrations used for capture were 0.5 μg/mL for PolySia and for HS. Association and dissociation phases were monitored during 200 s. All sensors were fully regenerated between experiments by dipping for 30 s in NaCl 2M solution.

### Sample preparation for label-free proteomics analysis

In order to analyze polySia carrier proteins, inflammatory monocytes were enriched from the bone marrow of WT and *St8sia4^-/-^*mice using a monocyte isolation kit as described previously (3 independent experiments, n=4 mice per group per experiment). WT polySia^low^ and polySia^high^ monocytes as well as St8sia4^-/-^ polySia^neg^ monocytes were then sorted by flow cytometry. Briefly, inflammatory monocytes were stained for 30 minutes at 4°C with fluorescent antibodies against CD11b, Ly6G, Ly6C, and polySia diluted in Cell Staining Buffer containing. Cell sorting was performed by flow cytometry using an Aria Fusion. Monocytes were identified as CD11b^+^/Ly6G^-^ /Ly6C^+^ cells with variable expression of polySia (PolySia^low^ and PolySia^high^ for WT cells and PolySia^neg^ for St8sia4^-/-^ cells). Cells were collected in 15 mL canonical tubes pre-coated with FBS, diluted with 10 mL DPBS and pelleted by centrifugation at 300 x g for 10 minutes. After a second washing with 10 mL DPBS, cell pellets were stored at - 80°C until use. Inflammatory monocytes pellets were lyzed in 250 mL of RIPA buffer (50 mM Tris-HCl pH8, 150 mM NaCl, 1% Nonidet P-40, 0.5% Sodium deoxycholate and 0.1% SDS) containing cOmplete™, EDTA-free Protease Inhibitor Cocktail (Sigma-Aldrich, #11873580001), 1 mg/mL DNAase I (Sigma-Aldrich #10104159001) and 5 mM Diisopropylfluorophosphate (Sigma; #D0879-1g). Cell lysates were then sonicated for 15 cycle (30 min ON / 30 min OFF) at 4°C and centrifuged at 20 000 x g for 15 minutes. Supernatants were transferred into new microtubes followed by measurement of protein concentration using DC protein assay kit II (Biorad; # 5000112). Immunoprecipitation (IP) experiment was performed using 40 μg of proteins of each sample diluted in 300 μL of RIPA buffer. Samples were first incubated with 12,5 μL of Protein G Dynabeads (ThermoFisher scientific; # 10004D) for 30 minutes at room temperature on a rotating wheel in order to remove non-specific binders. The supernatant was carefully separated from the magnetic beads using a magnetic rack (Bio-Rad; #1614916), transferred to a new microtube, and incubated with 2.5 μg of anti-polySia antibody (Absolute antibody; # Ab00240-23.0) overnight at 4°C on a rotating wheel. Samples were then incubated with 12.5 μL of Protein G Dynabeads (ThermoFisher scientific; # 10004D) for 3 hours at 4°C on a rotating wheel. The supernatant was removed using a magnetic rack, and the magnetic beads were subsequently washed 5 times with 300 μL of RIPA buffer. Finally, proteins retained on beads were eluted by adding 25 ml of 2X Laemmli buffer to the beads followed by incubation for 10 minutes at 60°c on a ThermoMixer. The eluted proteins were cleaned up on a short run on a 12% acrylamide SDS-PAGE gel followed by in gel trypsin digestion as described^82^ . Tryptic peptides were resuspended in 17 µL of 2% acetonitrile and 0.05% trifluoroacetic acid and analyzed by nano-liquid chromatography (LC) coupled to tandem MS, using an UltiMate 3000 system (NCS-3500RS Nano/Cap System; Thermo Fisher Scientific) coupled to an Orbitrap Exploris 480 mass spectrometer equipped with a FAIMS Pro device (Thermo Fisher Scientific). 4 µL of each sample was injected into the analytical C18 column (75 µm inner diameter × 50 cm, Acclaim PepMap 2µm C18 Thermo Fisher Scientific) equilibrated in 97.5% solvent A (5% acetonitrile, 0.2% formic acid) and 2.5% solvent B (80% acetonitrile, 0.2% formic acid). Peptides were eluted using a 2.5% - 40% gradient of solvent B over 122 min at a flow rate of 300 nl/min. The mass spectrometer was operated in data-dependent acquisition mode with the Xcalibur software. MS survey scans were acquired with a resolution of 60,000 and a normalized AGC target of 300 %. Two compensation voltages were applied (−45v /-60v). For 0.8s most intense ions were selected for fragmentation by high-energy collision-induced dissociation, and the resulting fragments were analyzed at a resolution of 15,000, using a standard AGC. Dynamic exclusion was used within 45s to prevent repetitive selection of the same peptide.

For cross-link experiment, BM inflammatory monocytes (5x10^5^ cells in 100 μL RPMI 1640 GlutaMAX medium / 1% PS / 10% FBS) from WT and *St8sia4^-/-^*mice (n= 4 per genotype) were plated in sterile V-bottom 96-well culture plate and incubated for 30 minutes at 4°C. CCL7-biotin (final concentration: 200 nM) was then added to the cells followed by another incubation for 30 minutes at 4°C. The 96-well culture plate was then centrifuged at 600 x g for 3 minutes at 4°C followed by removal of the supernatant by flicking the plate. Using a similar procedure, cells were washed twice with 200 μL of DPBS and finally resuspended in 150 μL of DPBS containing DTSSP (3,3’-dithiobis (sulfosuccinimidyl propionate, Thermo scientific, #21578) at either 0.25 and 1 mM for cross-linking validation by flow cytometry and 0.5 mM for proteomics analysis. Following incubation for 2 hours at 4°C, 50 μL of 100 mM Tris-HCl diluted in DPBS was added to stop the chemical reaction. The plate was then centrifuged at 600 x g for 3 minutes at 4°C followed by two washing of the cells with DPBS as described. For flow cytometry analysis, cells were labeled for 30 min at 4°C with 5 nM streptavidin-PE (#405204, BioLegend) diluted in FACS buffer (DPBS/ 2% FBS/ 2mM EDTA). For proteomics analysis, inflammatory monocytes pellets were lyzed in 100 mL of RIPA buffer (50 mM Tris-HCl pH8, 150 mM NaCl, 1% Nonidet P-40, 0.5% Sodium deoxycholate and 0.1% SDS) containing Halt™ Protease and Phosphatase Inhibitor Cocktail, EDTA-free (100X) (ThermoFischer Sceintific, #78441). Cell lysates were transferred into microtubes and diluted in 300 µL of DPBS containing Protease and Phosphatase Inhibitors. Cell lysates were then sonicated for 15 cycle (30 min ON / 30 min OFF) at 4°C and centrifuged at 20 000 x g for 10 minutes. Affinity purification was performed by incubating cell lysates with 50 µL of Streptavidine T1 Dynabeads™ MyOne™ (Invitrogen # 65601) suspension (pre-washed 4 times with DPBS according to manufacturer’s instructions) overnight at 4°C on a rotating wheel. The supernatant was carefully removed using a magnetic rack (Bio-Rad; #1614916), and the magnetic beads were washed three times with DPBS containing 0.01% Tween® 20 (Sigma-Aldrich; #P1379), followed by a final wash with DPBS alone. The beads were subsequently subjected to on-bead proteolysis. To that aim, beads were resuspended in 100 mM sodium bicarbonate buffer containing 4 % sodium deoxycholate (Sigma-Aldrich, #30970) and protein disulfide bonds were reduced and alkylated by addition of 10mM TCEP and 40mM chloroacetamide, at pH 8, 5min at 95°C. Proteins were digested using 1µg of trypsin, and resulting tryptic peptides were diluted with one volume of isopropanol containing 1% TFA, followed by desalting using SDB-RPS AttractSPE tips (Affinisep).

For proteome analysis of BM inflammatory monocytes, 1 × 10⁶ cells were plated in a V-bottom 96-well plate and lysed with 100 µL of 100 mM sodium bicarbonate, 4 % sodium deoxycholate, and heating 5min at 95°C. Half of the lysate volume was further processed by addition of 40 units of nuclease for DNA shearing, followed by protein reduction-alkylation, digestion with trypsin, and peptide desalting as described above. For mass spectrometry analysis, tryptic peptides were resuspended in DDM 0.015%, formic acid 0.1%, and an aliquot (50 ng) was analyzed by nanoLC-MS/MS using an UltiMate 3000 RS nanoLC system (ThermoFisher Scientific) coupled to a TIMS-TOF SCP mass spectrometer (Bruker). Peptides were separated on a C18 Aurora column (25 cm x 75 µm ID, IonOpticks) using a gradient ramping from 2% to 20% of B in 30 min, then to 37% of B in 3 min and to 85% of B in 2min (solvent A: 0.1% FA in H2O; solvent B 0.1% FA in acetonitrile), with a flow rate of 150 nl/min. MS acquisition was performed in DIA-PASEF mode on the precursor mass range [400-1000] m/z and ion mobility 1/K0 [0.64-1.37]. The acquisition scheme was composed of 8 consecutive TIMS ramps using an accumulation time of 100 ms, with 3 MS/MS acquisition windows of 25 Th for each of them. The resulting cycle time was 0.96 s. The collision energy was ramped linearly as a function of the ion mobility from 59 eV at 1/K0=1.6 Vs.cm−2 to 20 eV at 1/K0=0.6Vs.cm−2.

### Bioinformatics analysis of mass spectrometry raw files

Raw MS files from IP experiments were processed with the Mascot software (version 2.7.0) for database search and Proline for label-free quantitative analysis (version 2.1.2). Data were searched against *Mus musculus* entries of SwissProt 20201022 protein database (17,060 sequences). Carbamidomethylation of cysteines was set as a fixed modification, whereas oxidation of methionine and acetylation of proteins were set as variable modifications. Specificity of trypsin/P digestion was set for cleavage after K or R, and two missed trypsin cleavage sites were allowed. The mass tolerance was set to 5 ppm for the precursor and to 20 mmu in tandem MS mode. Minimum peptide length was set to 7 amino acids, and identification results were further validated in Proline by the target decoy approach using a reverse database at both a PSM and protein false-discovery rate of 1%. For label-free relative quantification of the proteins across biological replicates and conditions, cross-assignment of peptide ions peaks was enabled inside group with a match time window of 1 min, after alignment of the runs with a tolerance of +/- 600 s.

Median Ratio Fitting computes a matrix of abundance ratios calculated between any two runs from ion abundances for each protein. For each pair-wise ratio, the median of the ion ratios is then calculated and used to represent the protein ratio between these two runs. A least-squares regression is then performed to approximate the relative abundance of the protein in each run in the dataset. This abundance is finally rescaled to the sum of the ion abundances across runs.

Raw MS files from the cross-link and total proteome experiments were processed with Spectronaut 19.2 in directDIA+ mode. For the comparative proteome analysis of WT versus KO samples, data were searched against the Uniprot mouse reference proteome (downloaded April 2023). For the crosslinking experiment, searches were performed against the SwissProt mouse reference proteome (downloaded January 2024) supplemented with the human CCL7 protein sequence. In the Pulsar search, Carbamidomethyl (C) was specified as a fixed modification, and Acetyl (Protein N-term), Oxidation (M) as variable modifications. The peptide length range was set at 7–52, and 2 missed cleavages were allowed. For validation, the FDR was set at 1% at precursor, peptide and protein level. For Spectronaut DIA analysis, the Q-value cut-off at precursor level was at 0.01, and at protein level it was set respectively at 0.05 for each run and at 0.01 for the global dataset. Quantification of proteins was performed by calculating the median for each precursor of the intensity values retrieved for its MS2 fragment ions, and aggregating the different precursors abundances using the MaxLFQ algorithm. Global normalization was enabled.

Statistical analysis and volcano plots were performed using Microsoft Excel. A paired t-test was conducted on log₂-transformed values to assess differences in the abundance of proteins enriched by immunoprecipitation from FACS-sorted inflammatory monocytes. Significance level was set at p = 0.01, and ratios were considered relevant if higher than +/- 4. A two-sample equal variance t-test was performed on log2 transformed values to analyze differences in protein abundance in other biologic group comparisons. Significance level was set at p = 0.05, and ratios were considered relevant if higher than +/- 2.

The mass spectrometry proteomics data have been deposited to the ProteomeXchange Consortium via the PRIDE partner repository with the project accessions PXD070998 (related to figure 5C and table S2), PXD070693 (Figure 6D and table S3), and PXD070744 (Figure 7A and table S4).

### Adhesion assay

For the cell-adhesion assay, wells of a 96-well flat-bottom polystyrene high-bind microplate (Corning® #P3361) were left uncoated or coated overnight at 4 °C with one of the following in DPBS: 0.001% Poly-L-Lysine (Sigma-Aldrich #P4707), 12.5 µg/mL recombinant mouse ICAM-1/CD54 Fc chimera protein (Bio-Techne #796-IC), 100 µg/mL human fibronectin (Sigma-Aldrich #F2006), or 100 µg/mL human fibrinogen (Sigma-Aldrich #F4883). Purified inflammatory monocytes were then resuspended at 1×10^6^ cells/mL in RPMI 1640 GlutaMAX supplemented with 1% penicillin–streptomycin. After three additional washes of the wells with 200 µL DPBS, 1 × 10^5^ cells were added per well and incubated for 1 hour at 37 °C with 5% CO₂ to permit adhesion. Wells containing the cells were washed three times with 200 µL DPBS and cells were lysed with CellTiter-Glo 2.0 Reagent (#G9241; Promega) followed by luminescence measurement on a CLARIOstar. All assays were performed in quadruplicate. Results were expressed as the percentage of cells that have adhered relative to the initial inoculum.

### Immunofluorescence microscopy

Sterile glass coverslips (12 mm diameter; Fisher Scientific, #10006111) were placed in a 24-well cell culture plate, coated with 0.001% Poly-L-Lysine (Sigma-Aldrich, #P4707) for 1 hour at room temperature, and then washed five times with DPBS. Bone marrow-derived inflammatory monocytes (5 × 10^5^), purified as previously described previously, were resuspended in RPMI 1640 GlutaMAX medium supplemented with or without 10% FBS and 1% P/S, and incubated for 1 hour at 37 °C with 5% CO₂ to allow cell adhesion with or without spreading. CCR2 staining was performed on live cells after incubating the plate at 4 °C for 30 minutes followed three washes with 500 µL of cell staining buffer (CSB, BioLegend; #420201). Cells were incubated with 300 µL CSB containing TruStain FcX™ antibody solution (1:50 dilution; Biolegend; # 101320 for 30 minutes at 4 °C and then with 300 µL CSB containing TruStain FcX™ antibody solution and Alexa Fluor® 647 anti-mouse CD192 (CCR2) Antibody (1:200 dilution; Table S4). for 1 hour at 4 °C. Following three washes with 500 µL of cold sterile DPBS, cells were fixed with 4% paraformaldehyde (in sterile DPBS) for 15 minutes at room temperature. After fixation, cells were washed three times with 500 µL of cold sterile DPBS. To quench residual paraformaldehyde, cells were incubated with 100 mM glycine in DPBS for 5 minutes at room temperature. Regarding myosin and phalloidin staining, cells were fixed using paraformaldehyde as described above and then permeabilized for 10 minutes at room temperature using 0.1% Triton X-100. For microtubule staining, cells were fixed and permeabilized for 20 minutes at -20°C using cold methanol. In all conditions, cells were washed three times with 500 µL of cold sterile DPBS and non-specific binding sites were blocked by incubating the cells with 5% BSA in DPBS (filtered through a 0.2 µm membrane) for 1 hour at 4 °C. The blocking solution was then removed by aspiration, and 300 µL of blocking solution per well containing the primary anti-polySia, anti-MYH4, anti-MYH9 and anti-TUBA4A antibodies (Table S5) was added to each well followed by incubation overnight at 4 °C. The following day, cells were washed three times with 500 µL of cold sterile DPBS and incubated with 300 µL of blocking solution per well containing the secondary fluorescent anti-rabbit/mouse antibodies, fluorescent phalloidin and DAPI for 1 hour at room temperature (Table S5). After incubation, cells were washed three more times with 500 µL of cold sterile DPBS. Finally, coverslips were carefully removed and mounted onto glass slides using 10–15 µL of ProLong™ Glass Antifade Mountant (ThermoFisher Scientific; # P36984). Slides were allowed to cure overnight in the dark. Confocal microscopy images were acquired on a ZEISS LSM 880 equipped with a Plan-Apochromat 63×/1.4 oil objective. Data were collected in ZEN Black (v3.12) using frame scanning with line averaging = 2, 8-bit sampling, and a pinhole ∼1 Airy unit per channel. Fluorophores were recorded sequentially: DAPI (ex 405 nm, em 415–480 nm), Alexa Fluor 488 (ex 488 nm, em 493–555 nm), and Alexa Fluor 647 (ex 633 nm, em 638–755 nm). The pixel size was kept constant across all images (70.6 nm/pixel), and Z-stacks were acquired using the optimal step size automatically determined by ZEN (≈ 0.33 µm). For display, we used single optical sections (Figure 6) or maximum-intensity projections (Supplemental S) generated in either ZEN or Fiji: ImageJ. Scale bars were added in Fiji: ImageJ, and linear contrast adjustments were applied equally to all channels. Alternatively, confocal microscopy images were acquired on an Andor/Olympus Spinning Disk Confocal CSU-X1 equipped with a UPlanSApo 60x Oil Objective. Pixels resolution was measured at 0,135 µm and Z-stack were acquired with a step of 0,22µm. Acquisition was done sequentially with a camera IXon Ultra 888 EM 3000 preAMP gain 2, acquisition field was cropped to kept only homogeneous region. DAPI was acquired with laser line 405 at 5% (ILE 4 approx. 8.4mW) through a single band FF02-447/60-25 Filter (Semrock) and acquisition time was 150 ms. PolySia staining (Green) was acquired with laser line 488 at 7% (ILE 4 approx. 8,1mW) through a single band FF02-525/40-25 filter (Semrock) and acquisition time 50 ms. CCR2 staining (FarRed) was acquired with laser line 640 at 10% (ILE 4 approx. 15,3mW) through a single band FF02-685/40-25 filter (Semrock) and acquisition time 200 ms. The images were then processed in Fiji: ImageJ.

The two-colour three-dimensional random illumination microscopy (RIM) imaging was performed using the 3D RIM system.^71^ The homemade setup is coupled to an inverted microscope (TEi 2 Nikon) equipped with 60x magnification and a 1.30 N.A. objective (PLAN APO 60X ON 1.3 SIL λS NIKON). A pMOS camera (ORCA-Quest 2, C15550-22UP, Hamamatsu) is used for imaging. Diode lasers (Oxxius) with wavelengths centered at 488 nm (LBX-488-200-CSB) and 561 nm (LMX-561L-200-COL) are used for all experiments. Chroma’s 89901 ET - 405/488/561/640 nm Laser Quad Band Set multi-band excitation/multi-band emission filter has been used to separate the two colours. A binary spatial light modulator (QXGA fourth dimension), conjugated to the image plane and combined with polarization elements, is used to generate dynamic speckle on the object plane, as described.^71^ The synchronisation of the hardware (Z-platform, cameras, microscope, laser and SLM) is performed by an improved version of the commercial INSCOPER software.Fixed 3D RIM. Acquisition of the 40 image planes was performed sequentially with a 210 nm step. Two hundred speckles were used for each image plane to increase the desired super-resolved 3D resolution (110 nm, 110 nm, 300 nm). Image reconstruction was performed using the ALgoRIM software (https://github.com/teamRIM/tutoRIM). For the first channel images, the Wiener filter is set to 0.15, the deconvolution parameter to 0.08, and the regularization parameter to 0.15. For the second channel images, the Wiener filter is set to 0.25, the deconvolution parameter to 0.25 and the regularization parameter to 0.25. Any misalignment between the two colours and residual chromatic aberrations were corrected using SVI (Scientific Volume Imaging) software. The images were then processed in Fiji: ImageJ.

### Analysis of ST8SIA4 phenotypes in human

Phenotype association data (credible sets & colocalization results) were downloaded from the open target platform website (http://ftp.ebi.ac.uk/pub/databases/opentargets/platform/25.03/output/),^24^ and we considered all gwas traits displaying significant association with at least one SNP located less the 10 kb from *ST8SIA4* gene body for further analyses. When several studies reported associations for the same traits, we selected only the study with the most significant association. For the 3 phenotypes/studies selected for downstream analyses (GSCT90002340, GSCT90002316 and GSCT011096), we downloaded harmonized summary stats from the GWAS catalog^72^ (http://ftp.ebi.ac.uk/pub/databases/gwas/summary_statistics/) for plotting associations profiles. Haplotypes and LD analysis were performed with the LDlink website (https://ldlink.nih.gov/) using LDhap and LDmatrix functions. For colocalization analyses, we first considered colocalization results between the 3 focus traits and all eQTLs and sQTLs from the eQTL catalog^25^ and considered as significant any molecular QTL with a colocalization probability of > 0.8.

### Coverage QTL analyses

Coverage analyses were performed based on data from Quach et al., 2016 focusing on non-stimulated samples.^26^ Reads were realigned to hg38 genome using STAR^73^ and ggsashimi^74^ was used to compute coverage at base pair resolution in a window extending up to 10kb on each side of the *ST8SIA4* gene. For each individual, we then summed coverage values in bins of 2 kb, divided each coverage values by the total coverage across the full window. For each bin, we then applied an inverse-normal rank transform across individuals to remove the effects of outliers, and used Matrix eQTL to map coverage QTLs in a 100 kb window around the *ST8SIA4* gene, adjusting for age, % of European/African ancestry, GC content of the sample and sample-wide bias in relative coverage of 3’/5’ end of gene bodies. We then assessed significance of coverage QTLs, using a fastQTL-inspired permutation procedure.^75^ Briefly, we repeated the analyses on 100 permuted datasets, where genotypes labels were reassigned at random across individuals. For each coverage bin, we then extracted the smallest association p-value of each permutation to define the null distribution of association p-values in the absence of true signal. We then fitted a beta law to this distribution and used it to assign, for each bin, a p-value that the bin is under genetic control by at least one variant. We then applied a Benjamimini Hochberg correction across all 49 bins tested to adjust for the number of bind tested, and considered as significant any bin passing a 5% FDR threshold. Finally, for each significant bin, we finemapped associations using SuSiE,^76^ computing LD separately for European and African ancestry individual and using the average LD matrix for fine-mapping analysis. Finally, we tested each coverage bin with a significant QTL for colocalization with each *ST8SIA4*-associated phenotypes using the coloc.abf function from the coloc R package.^77^

### Quantification and statistical analysis

Where not specified otherwise, statistical analysis was performed with GraphPad Prism software (v10.6.1 and earlier versions). The statistical details can be found in figure legends. Experiments were analyzed using either unpaired or Welch’s t-tests (for populations with normal distribution with equal or unequal SD), Mann–Whitney tests (for populations without normal distribution) or paired *t*-tests. Correction for multiple comparison using Holm-Šídák method was applied when indicated. The normal distribution of the population was evaluated using D’Agostino-Pearson omnibus normality test. Results are expressed as mean ± standard deviation (SD) or ± standard error of the mean (SEM). A *p*-value ≤0.05 was considered significant, with *, **, ***, **** indicating (adjusted) *p*-values ≤0.05, ≤0.01, ≤0.001 and ≤0.0001 respectively. The abbreviation ’ns’ stands for non-significant. Statistical analysis of proteomics data was performed using Microsoft Excel (Version 16.89 and earlier versions). A paired t-test was conducted on log₂-transformed values to assess differences in the abundance of proteins enriched by immunoprecipitation from FACS-sorted inflammatory monocytes. A two-sample equal variance t-test was performed on log2 transformed values to analyze differences in protein abundance in other biologic group comparisons.

