## Supplementary figures for "ST8SIA4-mediated polysialylation is critical for CCR2-driven monocyte egress from the bone marrow"

### SUPLEMENTARY DATA

**SUPLEMENTARY FIGURES**

Figure S1: Genetic architecture of the ST8SIA4 locus and its association with circulating monocyte and lymphocyte counts, and systemic lupus erythematosus (related to figure 1 and table S1).

Figure S2: Gating strategies used for flow cytometry analysis of blood and lung immune cells (related to figure 2).

Figure S3: Flow cytometry analysis of lung immune cells in mice infected with *Mycobacterium tuberculosis*. (related to figure 2).

Figure S4: Analysis of inflammatory monocyte development in WT and *St8sia4*$^{-/-}$ mice (related to figure 3)

Figure S5: Analysis of monocyte phenotype in WT and *St8sia4*$^{-/-}$ mice (related to figures 3 and 4)

Figure S6: Analysis of the role of NCAM1, polySia and glycosaminoglycans in chemokine binding (related to figures 5-6).

Figure S7: *St8sia4* deficiency alters CCR2 surface localization and function as well as cytoskeletal architecture (related to figures 6-7 and tables S3-4)

**SUPLEMENTARY TABLES**

(in separate files)

Table S1: Functional genetic analysis of the association between *ST8SIA4* variants and circulating monocyte and lymphocyte counts, and systemic lupus erythematosus

Table S2: Proteomics analysis of proteins enriched by anti-polySia immunoprecipation from bone marrow polySia$^{Low}$ and polySia$^{High}$ WT inflammatory monocytes compared to *St8sia4*$^{-/-}$ inflammatory monocytes (related to figure 6).

Table S3: Proteomics analysis of proteins enriched following CCL7-crosslinking on bone marrow WT inflammatory monocytes compared to *St8sia4*$^{-/-}$ inflammatory monocytes (related to figure 7).

Table S4: Comparative proteomics analysis of proteins from bone marrow WT inflammatory monocytes and *St8sia4*$^{-/-}$ inflammatory monocytes (related to Figure 6).

Table S5: List of the antibodies used for flow cytometry and microscopy in the study

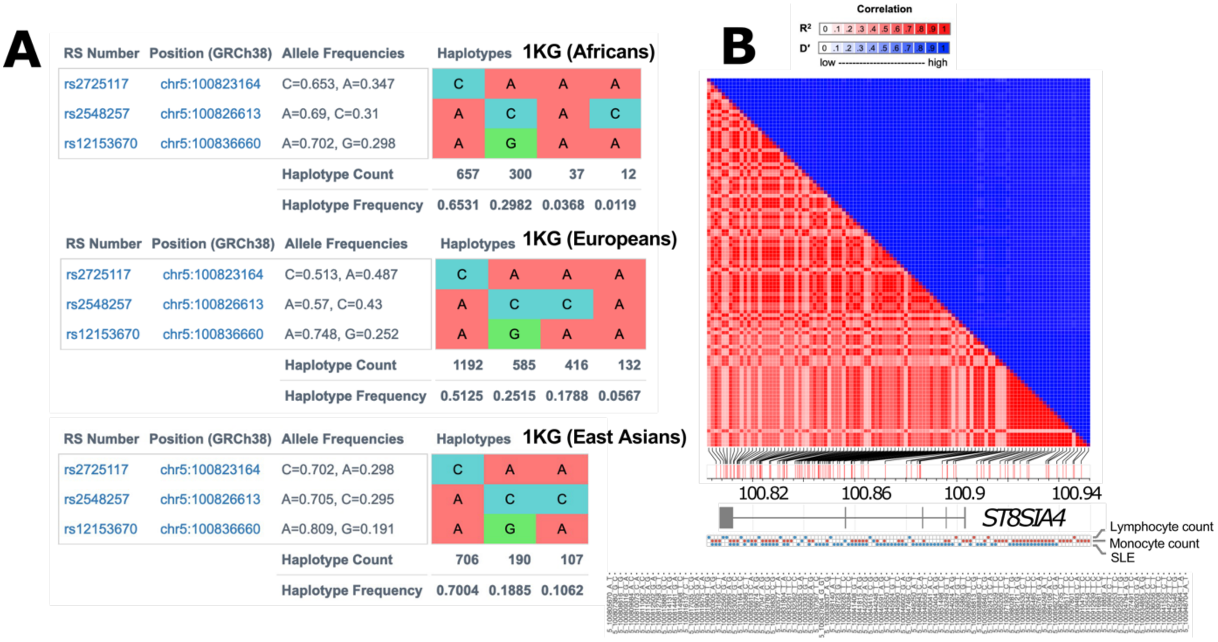

A

**1KG (Africans)**

| RS Number | Position (GRCh38) | Allele Frequencies | Haplotypes | | | |
|---|---|---|---|---|---|---|
| rs2725117 | chr5:100823164 | C=0.653, A=0.347 | C | A | A | A |
| rs2548257 | chr5:100826613 | A=0.69, C=0.31 | A | C | A | C |
| rs12153670 | chr5:100836660 | A=0.702, G=0.298 | A | G | A | A |
| | | Haplotype Count | 657 | 300 | 37 | 12 |
| | | Haplotype Frequency | 0.6531 | 0.2982 | 0.0368 | 0.0119 |

**1KG (Europeans)**

| RS Number | Position (GRCh38) | Allele Frequencies | Haplotypes | | | |
|---|---|---|---|---|---|---|
| rs2725117 | chr5:100823164 | C=0.513, A=0.487 | C | A | A | A |
| rs2548257 | chr5:100826613 | A=0.57, C=0.43 | A | C | C | A |
| rs12153670 | chr5:100836660 | A=0.748, G=0.252 | A | G | A | A |
| | | Haplotype Count | 1192 | 585 | 416 | 132 |
| | | Haplotype Frequency | 0.5125 | 0.2515 | 0.1788 | 0.0567 |

**1KG (East Asians)**

| RS Number | Position (GRCh38) | Allele Frequencies | Haplotypes | | |
|---|---|---|---|---|---|
| rs2725117 | chr5:100823164 | C=0.702, A=0.298 | C | A | A |
| rs2548257 | chr5:100826613 | A=0.705, C=0.295 | A | C | C |
| rs12153670 | chr5:100836660 | A=0.809, G=0.191 | A | G | A |
| | | Haplotype Count | 706 | 190 | 107 |
| | | Haplotype Frequency | 0.7004 | 0.1885 | 0.1062 |



**Figure S1: Genetic architecture of the ST8SIA4 locus and its association with monocyte and lymphocyte counts and systemic lupus erythematosus (related to figure 1).**

(A) Allelic and haplotypic frequencies in humans for the three index single-nucleotide polymorphisms (SNPs) associated with monocyte and lymphocyte counts and systemic lupus erythematosus.

(B) Linkage-disequilibrium (LD) structure between 107 phenotype-associated genetic variants at the *ST8SIA4* locus. The lower panel displays 95% credible sets and the direction of association for each phenotype (red = increased; blue = decreased phenotype among carriers of the alternative allele)

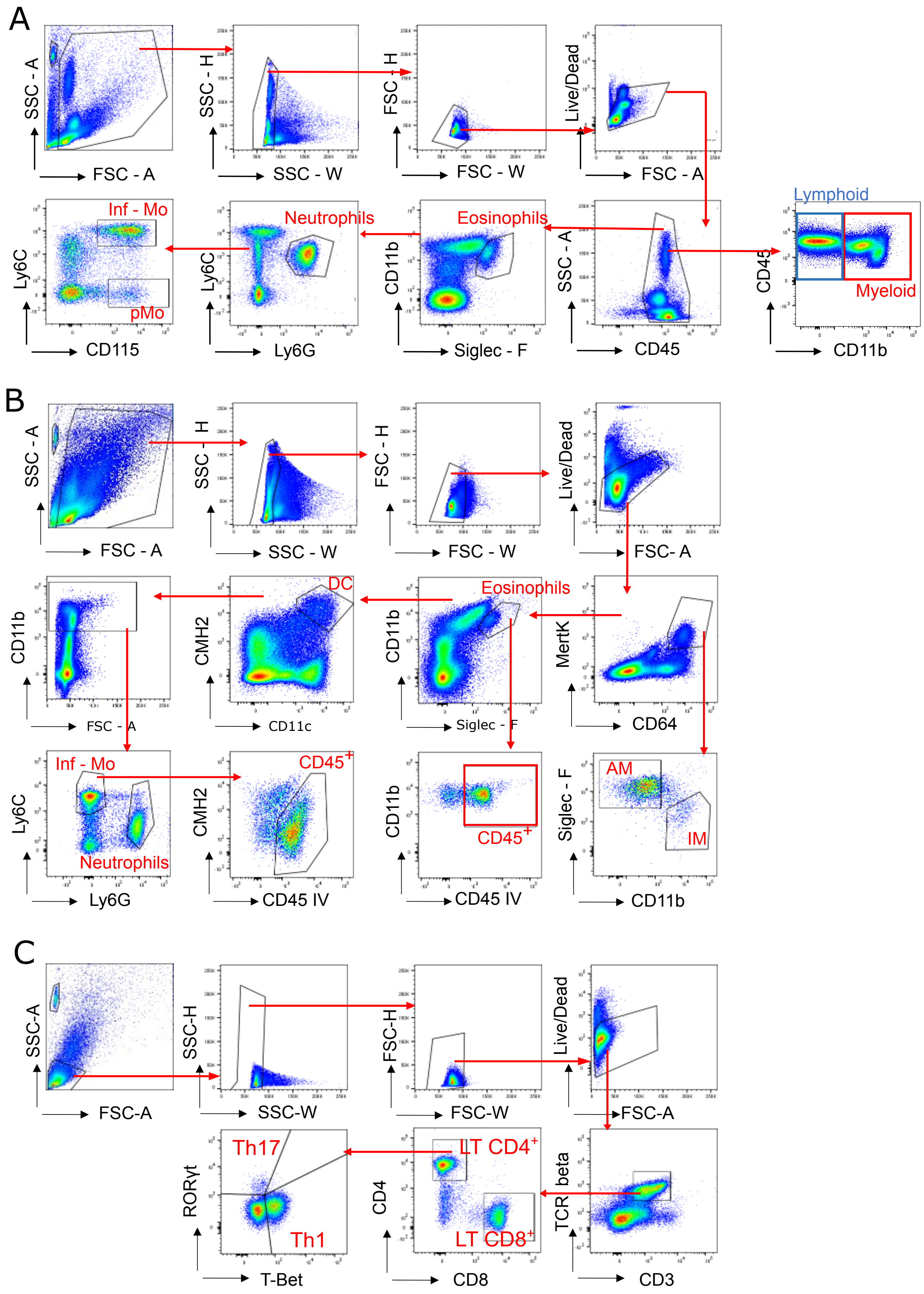

A
SSC - A
FSC - A
SSC - H
SSC - W
FSC - H
FSC - W
Live/Dead
FSC - A
Lymphoid
Myeloid
CD45
CD11b
SSC - A
CD45
Eosinophils
CD11b
Siglec - F
Neutrophils
Ly6C
Ly6G
Inf - Mo
pMo
Ly6C
CD115
B
SSC - A
FSC - A
SSC - H
SSC - W
FSC - H
FSC - W
Live/Dead
FSC- A
MertK
CD64
Eosinophils
CD11b
Siglec - F
DC
CMH2
CD11c
CD11b
FSC - A
Inf - Mo
Neutrophils
Ly6C
Ly6G
CD45+
CMH2
CD45 IV
CD45+
CD11b
CD45 IV
AM
IM
Siglec - F
CD11b
C
SSC-A
FSC-A
SSC-H
SSC-W
FSC-H
FSC-W
Live/Dead
FSC-A
TCR beta
CD3
LT CD4+
LT CD8+
CD4
CD8
Th17
Th1
RORγt
T-Bet

**Figure S2: Gating strategies used for flow cytometry analysis of blood and lung immune cells (related to figure 2)**

(A) Gating strategy used for flow cytometry analysis of mouse circulating lymphoid ($CD45^+$ $CD11b^-$) and myeloid cells ($CD45^+$ $CD11b^+$) as well as myeloid cell subtypes in mice. Inf-Mo and pMo stand for inflammatory and patrolling monocytes respectively.

(B) Gating strategy used for flow cytometry analysis of pulmonary myeloid cell subtypes in mice infected with Mtb. AM and IM stand for alveolar and interstitial macrophages respectively. DC stand for dendritic cells. CD45.2 labeling of eosinophils and inflammatory monocytes (inf-Mo) is shown as a representative example to distinguish their localization in the lung vasculature versus the tissue parenchyma.

(C) Similar to panel B for $CD4^+$ and $CD8^+$ lymphocytes (LT) subpopulations including T-helper (Th) 1 and 17.

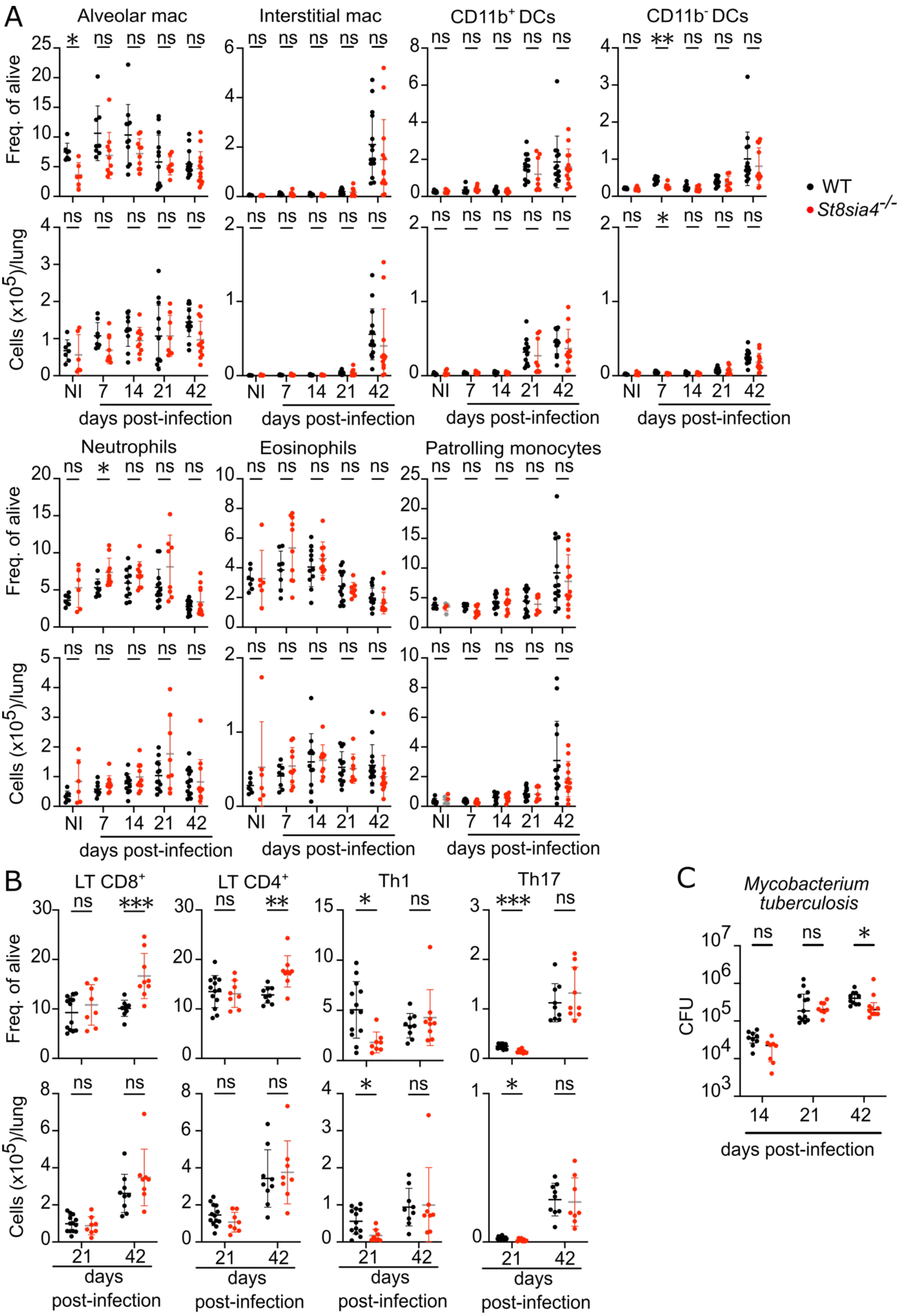

A
Alveolar mac
Interstitial mac
CD11b+ DCs
CD11b- DCs
Freq. of alive
Cells (x10^5)/lung
NI 7 14 21 42
days post-infection
WT
St8sia4-/-
Neutrophils
Eosinophils
Patrolling monocytes
B
LT CD8+
LT CD4+
Th1
Th17
days post-infection
C
Mycobacterium tuberculosis
CFU
14 21 42
days post-infection

**Figure S3: Flow cytometry analysis of lung immune cells in mice infected with *Mycobacterium tuberculosis* (related to figure 2).**

(A) Flow cytometry measurement of the frequency (top panel) and number (bottom panel) of myeloid cell subtypes in non-infected (NI) and during infection of WT and *St8sia4*$^{-/-}$ mice with *Mycobacterium tuberculosis*. Each data point corresponds to an individual mouse (n=6-14 mice per group per time point, pooled from at least 2 independent experiments per time point). Bars and error indicate the mean ± standard deviation. Statistical analysis was performed using Mann-Whitney *t*-tests corrected for multiple comparison using Holm-Šídák method. $*$ and $**$ indicate adjusted *p*-values ≤0.05 and ≤0.01 respectively. The abbreviation 'ns' stands for non-significant.

(B) Similar to panel B for CD4$^+$ and CD8$^+$ lymphocytes (LT) as well as T-helper (Th) 1 (CD4$^+$ T-bet$^+$ lymphocytes) and Th17 cells (CD4$^+$ ROR$\gamma$t$^+$ lymphocytes). Each data point corresponds to an individual mouse (n=8-13 mice per group per time point, pooled from at least 2 independent experiments). Bars and error indicate the mean ± standard deviation. Statistical analysis was performed using Mann-Whitney *t*-tests corrected for multiple comparison using Holm-Šídák method. $*$, $**$ and $***$ indicate adjusted *p*-values ≤0.05, ≤0.01 and ≤0.001 respectively. The abbreviation 'ns' stands for non-significant.

(C) Colony forming units (CFU) of *Mycobacterium tuberculosis* in the lung of WT and *St8sia4*$^{-/-}$ mice at 14-, 21- and 42-days post-infection. Each data point corresponds to an individual mouse (n=8-13 mice per group per time point, pooled from at least 2 independent experiments). Bars and error indicate the median ± interquartile range. Statistical analysis was performed using Mann-Whitney *t*-tests corrected for multiple comparison using Holm-Šídák method. $*$ indicates an adjusted *p*-values ≤0.05, whereas the abbreviation 'ns' stands for non-significant.

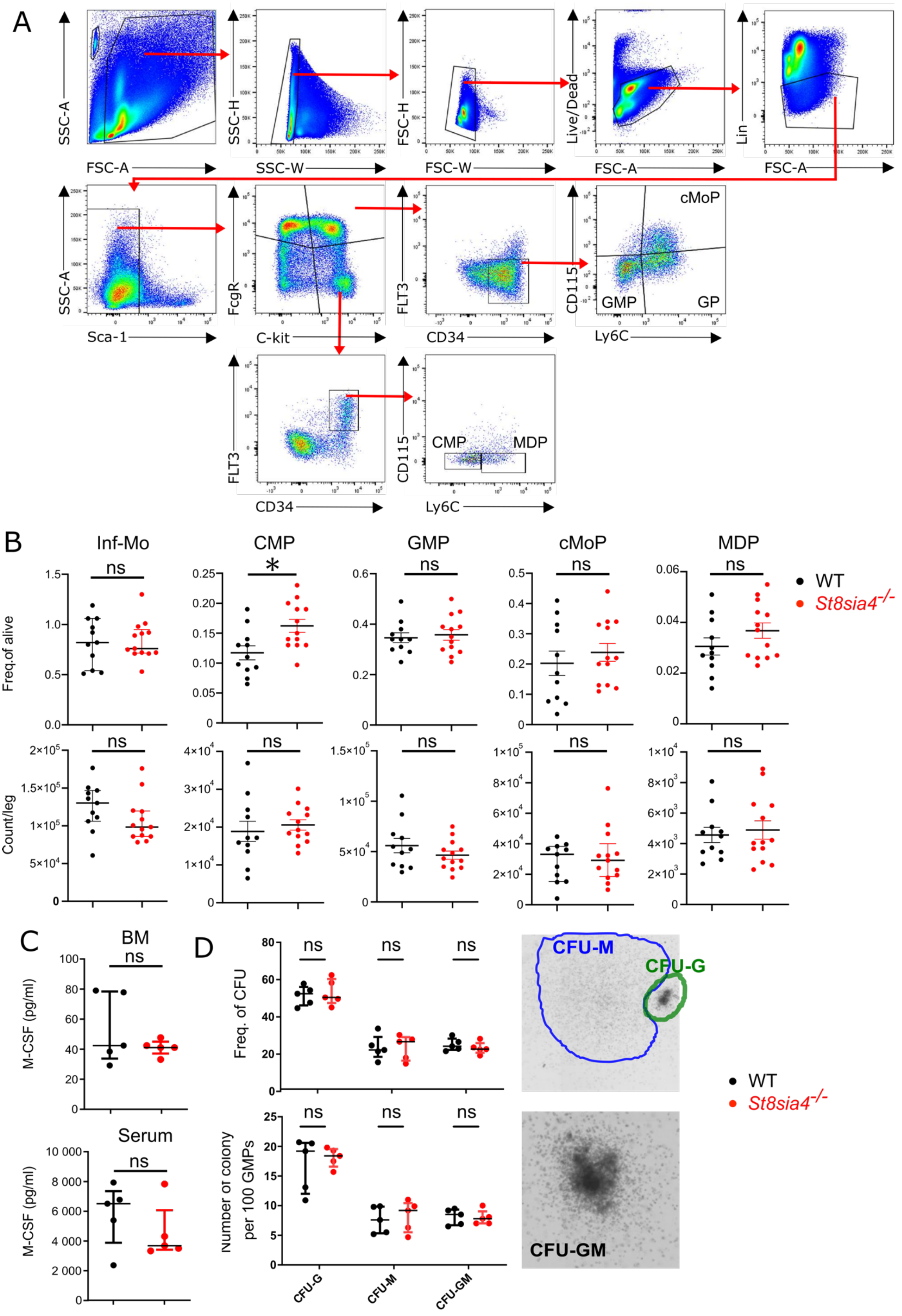
A
SSC-A
FSC-A
SSC-H
SSC-W
FSC-H
FSC-W
Live/Dead
FSC-A
Lin
FSC-A
SSC-A
Sca-1
FcgR
C-kit
FLT3
CD34
CD115
Ly6C
cMoP
GMP
GP
FLT3
CD34
CD115
Ly6C
CMP
MDP
B
Inf-Mo
CMP
GMP
cMoP
MDP
Freq.of alive
Count/leg
ns
*
WT
St8sia4-/-
C
BM
Serum
M-CSF (pg/ml)
D
Freq. of CFU
Number of colony per 100 GMPs
CFU-G
CFU-M
CFU-GM
WT
St8sia4-/-

**Figure S4: Analysis of inflammatory monocyte development in WT and *St8sia4*$^{-/-}$ mice (related to figure 3).**

(A) Gating strategy used for flow cytometry analysis of hematopoietic cell subpopulations in the bone marrow (BM) of WT and *St8sia4*$^{-/-}$ mice.

(B) Flow cytometry measurement of the frequency (top panel) and number (bottom panel) of inflammatory monocytes (Inf-Mo) and their hematopoietic progenitors in the BM of WT and *St8sia4*$^{-/-}$ mice. CMP: multipotent common myeloid progenitor; GMP: granulocyte-monocyte progenitors; cMoP: Common Monocyte Progenitors; MDP: Monocyte-DC Progenitors. Each data point corresponds to an individual mouse (n=11-13 mice per group, pooled from 4 independent experiments). Bars and error indicate the mean ± standard error of the mean. Statistical analysis was performed using Mann-Whitney *t*-tests. * indicates a *p*-value ≤0.05. The abbreviation 'ns' stands for non-significant.

(C) Concentration of Macrophage Colony Stimulating Factor (M-CSF) in the BM supernatant (top panel) and serum (bottom panel) of WT and *St8sia4*$^{-/-}$ mice, as detected by flow cytometry using a multiplex bead-based assay. Each data point corresponds to an individual mouse (n=5 mice per group). Bars and error indicate the median ± interquartile range. Statistical analysis was performed using Mann-Whitney *t*-tests. The abbreviation 'ns' stands for non-significant.

(D) Frequency (top left), total number (bottom left), and morphology (right panels) of granulocyte (G), macrophage (M), and granulocyte-macrophage (GM) colonies generated in a colony formation assay using FACS-sorted GMPs isolated from the bone marrow of WT and *St8sia4*$^{-/-}$ mice. Statistical analysis was performed using Mann-Whitney *t*-tests corrected for multiple comparison using Holm-Šídák method. The abbreviation 'ns' stands for non-significant.

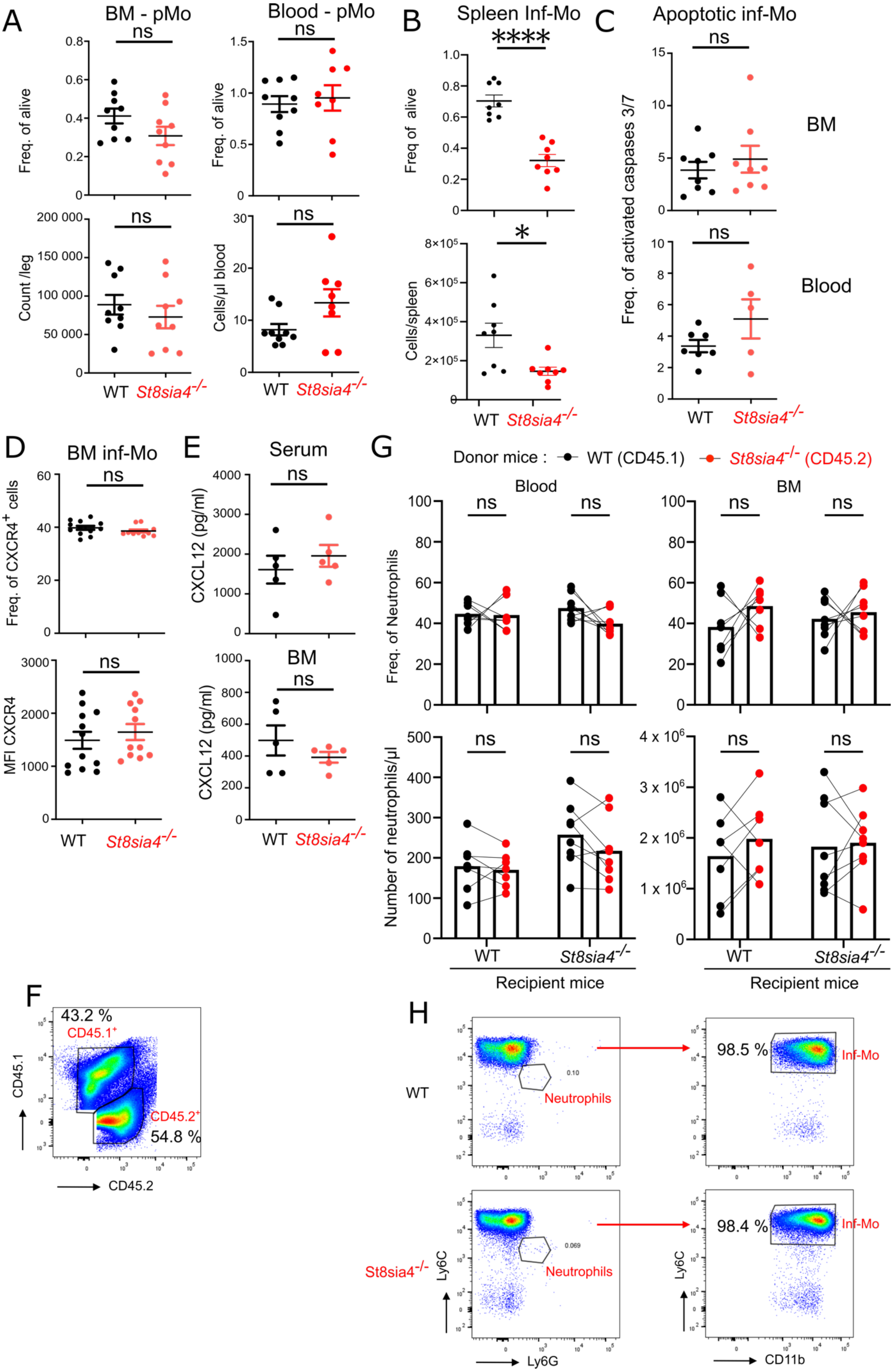

A
BM - pMo
Blood - pMo
Freq. of alive
Count /leg
Cells/µl blood
WT
St8sia4-/-
B
Spleen Inf-Mo
Freq of alive
Cells/spleen
C
Apoptotic inf-Mo
Freq. of activated caspases 3/7
BM
Blood
D
BM inf-Mo
Freq. of CXCR4+ cells
MFI CXCR4
E
Serum
BM
CXCL12 (pg/ml)
G
Donor mice : WT (CD45.1) St8sia4-/- (CD45.2)
Blood
BM
Freq. of Neutrophils
Number of neutrophils/µl
Recipient mice
F
43.2 %
CD45.1+
CD45.2+
54.8 %
CD45.1
CD45.2
H
WT
St8sia4-/-
Neutrophils
98.5 %
98.4 %
Inf-Mo
Ly6C
Ly6G
CD11b

**Figure S5: Analysis of monocyte phenotype in WT and *St8sia4$^{-/-}$* mice (related to figures 3 and 4).**

(A) Flow cytometry analysis of the frequency (top) and number (bottom) of bone marrow (BM) and blood patrolling monocytes (pMo) in WT and *St8sia4$^{-/-}$* mice. Each data point corresponds to an individual mouse (n= 8-9 mice per group, pooled from 3 independent experiments). Bars and error indicate the mean ± standard error of the mean. Statistical analysis was performed using Wech's *t*-tests. The abbreviation 'ns' stands for non-significant.

(B) Frequency (top) and number (bottom) of splenic inflammatory monocytes (inf-Mo) in WT and *St8sia4$^{-/-}$* mice as measured by flow cytometry. Each data point corresponds to an individual mouse (n=8 mice per group, pooled from 3 independent experiments). Bars and error indicate the mean ± standard error of the mean. Statistical analysis was performed using Wech's *t*-tests. ∗ and ∗∗∗∗ indicate *p*-values ≤0.05 and ≤0.0001.

(C) Flow cytometry analysis of the frequency of apoptotic (activated caspase 3/7$^{+}$) inflammatory monocytes (inf-Mo) in bone marrow (top) and blood (bottom) of WT and *St8sia4$^{-/-}$* mice. Each data point corresponds to an individual mouse (n= 5-8 mice per group, pooled from 2 independent experiments). Bars and error indicate the mean ± standard error of the mean. Statistical analysis was performed using Mann-Whitney *t*-tests. The abbreviation 'ns' stands for non-significant.

(D) Frequency (top) and representative median fluorescence intensity (MFI; bottom panel) of CXCR4 surface expression on BM inflammatory monocytes from WT and *St8sia4$^{-/-}$* mice, as detected by flow cytometry. Each data point corresponds to an individual mouse (n= 11-12 mice per group, pooled from 2 independent experiments). Bars and error indicate the mean ± standard error of the mean. Statistical analysis was performed using Wech's *t*-tests. The abbreviation 'ns' stands for non-significant.

(E) Serum (top) and BM extracellular fluid (bottom) CXCL12 concentration in WT and *St8sia4$^{-/-}$* mice, as measured by flow cytometry. Each data point corresponds to an individual mouse (n= 5 mice per group). Bars and error indicate the mean ± standard error of the mean. Statistical analysis was performed using Mann-Whitney *t*-tests. The abbreviation 'ns' stands for non-significant.

(F) Representative FACS plot showing the BM cell mixture from WT (CD45.1$^{+}$) and *St8sia4$^{-/-}$* (CD45.2$^{+}$) mice used for generating of mixed BM chimera.

(G) Flow cytometry analysis of the proportion (top) and number (bottom) of neutrophils derived from WT (CD45.1$^{+}$) and *St8sia4$^{-/-}$* (CD45.2$^{+}$) donor mice in the blood (left panel) and BM (right panel) from irradiated WT and *St8sia4*$^{-/-}$ recipient mice. Each data point represents either donor CD45.1$^{+}$ WT cells (black) or CD45.2$^{+}$ ST8SIA4-deficient cells (red) (n=8 mice per group, pooled from 2 independent experiments). Connecting lines indicate that paired data originate from the same recipient mouse, while the bars represent the mean value across all mice Statistical analysis was performed using paired *t*-tests corrected for multiple comparison using Holm-Šídák method. The abbreviation 'ns' stands for non-significant.

(H) Representative flow cytometry plots showing the purity of BM inf-Mo isolated from WT and *St8sia4*$^{-/-}$ mice.

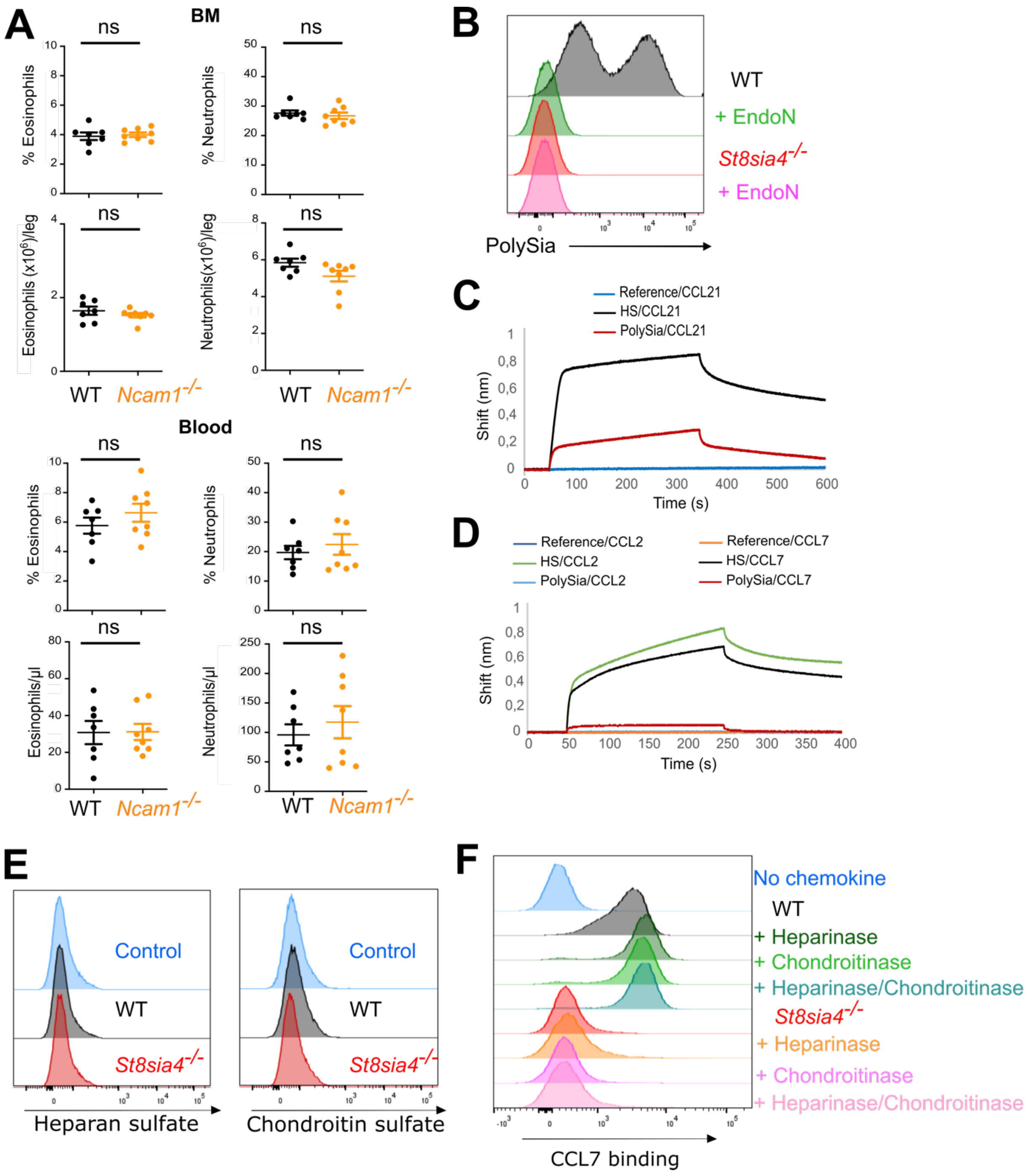


**Figure S6: Analysis of the role of NCAM1, polySia and glycosaminoglycans in chemokine binding (related to figures 5-6).**

(A) Flow cytometry measurement of the frequency (top) and number (bottom) of bone marrow (BM) and blood eosinophils (left) and neutrophils (right) in WT and *Ncam1*$^{-/-}$ mice. Each data point corresponds to an individual mouse (n= 7-8 mice per group). Bars and error indicate the mean ± standard error of the mean. Statistical analysis was performed using Mann-Whitney *t*-tests. The abbreviation 'ns' stands for non-significant.

(B) Representative flow cytometry histograms of cell surface polySia on WT and *St8sia4*$^{-/-}$ inflammatory monocytes treated or not with Endo-N-acylneuraminidase (endoN). (n=3 independent experiment with pooled monocytes purified from 3-4 mice per group).

(C) Biolayer Interferometry (BLI) sensorgrams of human CCL21 (200 nM) binding to polySia and heparan sulfate (HS)**.**

(D) Similar to panel C for human CCL2 and CCL7 (1 $\mu$M each).

(E) Representative flow cytometry histograms of cell surface heparan sulfate and chondroitin sulfate on WT and *St8sia4*$^{-/-}$ BM inflammatory monocytes (n=2 independent experiment with pooled monocytes purified from 3-4 mice per group).

(F) Flow cytometry histograms of the binding of biotinylated CCL7, coupled to fluorescent streptavidin, to WT and *St8sia4*$^{-/-}$ BM inflammatory monocytes prior to or after treatment of the cells with heparinases and/or chondroitinases (n=1 experiment with pooled monocytes purified from 3-4 mice per group).

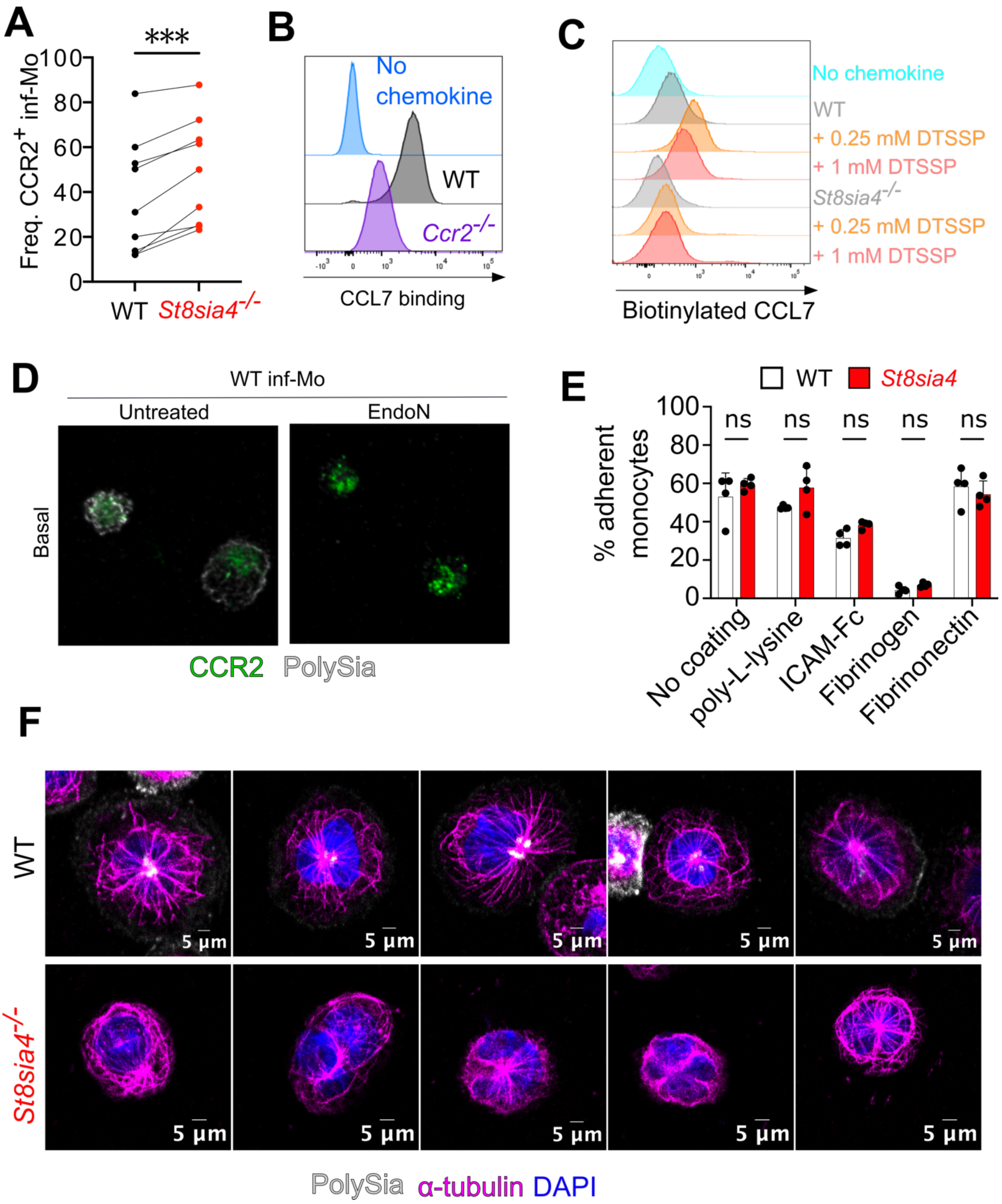


**Figure S7: St8sia4 deficiency alters CCR2 surface localization and function as well as cytoskeletal architecture (related to figures 6-7 and table S4)**

(A) Frequency of CCR2+ inflammatory monocytes (Inf-Mo) purified from the bone marrow of WT and *St8sia4-/-* mice, as measured by flow cytometry. Each data point corresponds to an individual experiment (n=9 experiments with pooled BM monocytes from 3-4 mice per group per experiment). Bars represent the mean, while lines connect paired WT and ST8SIA4-deficient cells from the same experiment. Statistical analysis was performed using a paired *t*-test. ∗∗∗ indicates a *p*-values ≤0.001.

(B) Flow cytometry histograms of the binding of biotinylated CCL7, coupled to fluorescent streptavidin, to WT and *Ccr2*$^{-/-}$ BM inflammatory monocytes (n=1 experiment with pooled monocytes purified from 4 mice per group).

(C) Flow cytometry histograms showing the presence of biotinylated CCL7 on the cell surface of WT and *St8sia4*$^{-/-}$ BM inflammatory monocytes that were treated or not with two concentrations of the protein crosslinking reagent DTSSP (3,3'-dithiobis(sulfosuccinimidyl propionate)). In absence of cross-linker, biotinylated CCL7 is almost completely removed from cell surface of WT cells due to the numerous washing step of the procedure (n=1 experiment with pooled monocytes purified from 4 mice per group).

(D) Confocal microscope images of cell surface CCR2 (in green) and polySia (in white) in WT BM inflammatory monocytes, treated or not with EndoN. Z-stack images representing the basal, medial, and apical sections of the cells are shown. Images were processed using Fiji: ImageJ software and are representative of a single optical section representing the basal region of the cells.

(E) Percentages of WT and *St8sia4*$^{-/-}$ inflammatory monocytes adhering to wells coated with different substrates. Percentages were calculated as the proportion of cells remaining after 1 hour of culture and three washing steps, relative to the initial number of cells seeded. Cell quantification was performed using the CellTiter-Glo® 2.0 Assay. Each data point corresponds to an individual mouse (n=4 mice per group). Bars and error indicate the mean ± standard deviation. Statistical analysis was performed using Wech's *t*-tests corrected for multiple comparison using Holm-Šídák method. The abbreviation 'ns' stands for non-significant.

(F) Confocal microscope images of microtubule network ($\alpha$-tubulin in magenta), polySia (in white), and cell nucleus (DAPI in blue) in WT and *St8sia4*$^{-/-}$ inflammatory monocytes. Images were processed using Fiji: ImageJ software and are representative of a single optical section above the basal region of the cells.
